# Dopamine Promotes Cognitive Effort by Biasing the Benefits Versus Costs of Cognitive Work

**DOI:** 10.1101/778134

**Authors:** A. Westbrook, R. van den Bosch, J. I. Määttä, L. Hofmans, D. Papadopetraki, R. Cools, M. J. Frank

## Abstract

Stimulants like methylphenidate are increasingly used for cognitive enhancement, but precise mechanisms are unknown. We found that methylphenidate boosts willingness to expend cognitive effort by altering the benefit-to-cost ratio of cognitive work. Willingness to expend effort was greater for participants with higher striatal dopamine synthesis capacity, while methylphenidate and sulpiride – a selective D2 receptor antagonist – increased cognitive motivation more for participants with lower synthesis capacity. A sequential sampling model informed by momentary gaze revealed that decisions to expend effort are related to amplification of benefit-versus-cost information attended early in the decision process, while the effect of benefits is strengthened with higher synthesis capacity and by methylphenidate. These findings demonstrate that methylphenidate boosts the perceived benefits-versus-costs of cognitive effort by modulating striatal dopamine signaling.

**One Sentence Summary:** Striatal dopamine increases cognitive effort by respectively amplifying and attenuating the subjective benefits and costs of cognitive control.

---

Cognitive control is effortful, causing people to avoid demanding tasks (*1*) and to discount goals (*2, 3*). Striatal dopamine invigorates physical action by mediating cost-benefit tradeoffs (*4*). In cortico-striatal loops, dopamine has opponent effects on D1 and D2-expressing medium spiny neurons, which modulate sensitivity to the benefits versus the costs of actions (*5*). Given that similar mechanisms may govern cognitive action selection (*6-8*), we hypothesized that striatal dopamine could promote willingness to exert cognitive effort, enhancing attention, planning, and decision-making (*8-11*).

Converging evidence on cognitive motivation in Parkinson’s disease (*12-15*) provides an initial basis for this conjecture. Moreover, catecholamine-enhancing psychostimulants alter cognitive effort in rodents (*10*) and humans (*16*). This raises the question of whether so called smart drugs act by enhancing the willingness rather than the ability to exert cognitive control. Indeed, the dominant interpretation is that stimulants improve cognitive processing, via direct cortical effects, noradrenaline transmission (*17, 18*) and/or concomitant working memory improvements (*19*). We instead hypothesized that methylphenidate (a dopamine and noradrenaline reuptake blocker) boosts cognitive control by increasing striatal dopamine and, accordingly, sensitivity to the benefits-versus-costs of cognitive effort.

50 healthy, young adults (ages 18—43, 25 men) completed a cognitive effort discounting paradigm (*2*) quantifying the subjective effort costs as the amount of money required to make participants equally willing to perform a hard (N = 2, 3, 4) versus easier (N = 1, 2) level of the N-back working memory task. We defined the subjective value of an offer to complete a harder task (N = 2—4) as the amount offered for the task, minus subjective costs.

Subjective values decreased with N-back load, indicating rising subjective costs (Fig. 1A). Critically, greater willingness to expend cognitive effort corelated with higher dopamine synthesis capacity (measured using [^18^F]DOPA PET) in the caudate nucleus (independently defined (*20*); Fig. 1A—C, Fig. S1). A mixed-effects model confirmed that on placebo, subjective values increased with larger offer amounts (€4 versus €2 offers; *β* = 0.022, *P* = 0.011), smaller relative load (*β* = −0.15, *P* = 8.9×10^−15^), and higher dopamine synthesis capacity (*β* = 0.064, *P* = 0.022). These individual difference effects were selective to the caudate nucleus (Fig. S1—S2), consistent with human imaging studies on cognitive motivation (*7, 21, 22*). Although N-back performance decreased with load, dopamine effects on discounting could not be attributed to performance changes (Supplemental Results). Moreover, there were no drug effects on performance because drugs were administered after N-back.

**Fig. 1.**
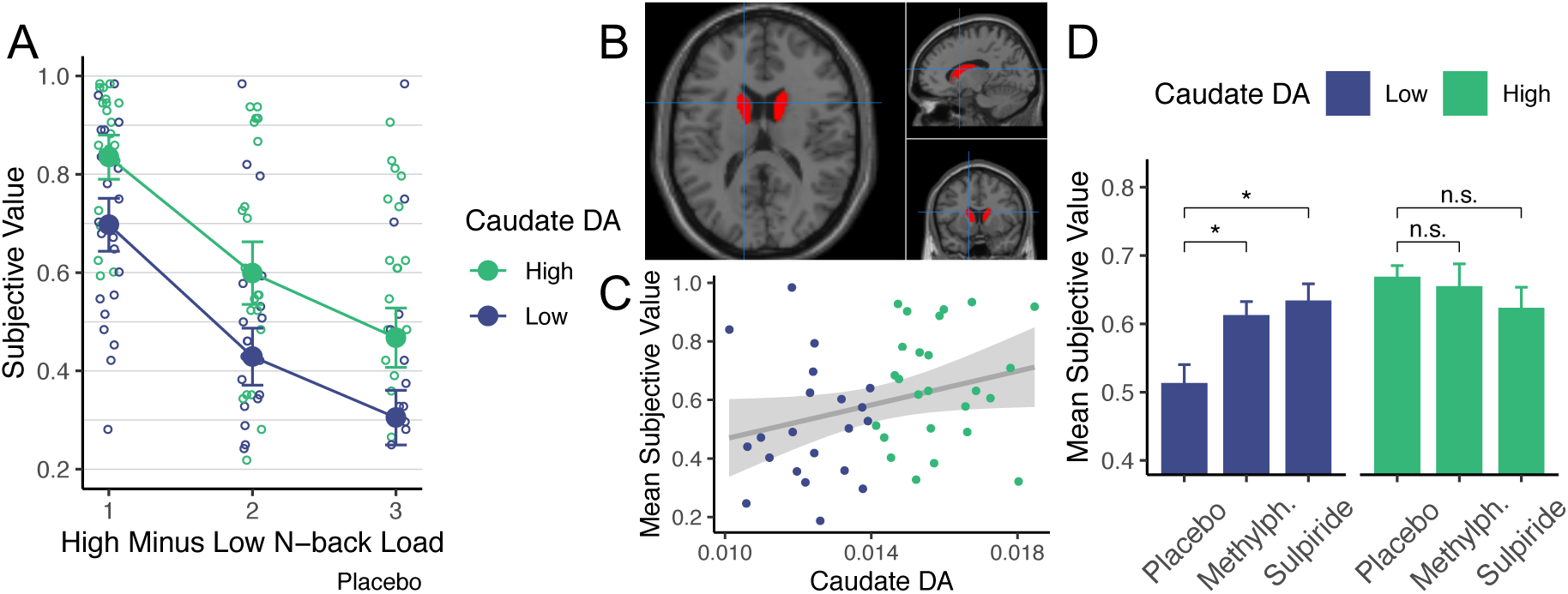
Participants discounted offers as a function of cognitive load, dopamine synthesis capacity (DA), and drug. **A.** Offers were discounted more for high-versus low-load levels, and more by participants with below-versus above-median dopamine synthesis capacity. Circles show individual’s indifference points. Filled circles show group mean +/- SEM. **B.** Caudate nucleus mask. Crosshairs at MNI [-14, 10, 16]. **C.** Participant-averaged subjective values correlated with synthesis capacity on placebo (Spearman *r* = 0.32, *P* = 0.029). **D.** Methylphenidate (*t*_*paired*_(22) = 2.29, *P* = 0.032) and sulpiride (*t*_*paired*_(22) = 2.36, *P* = 0.028) increased subjective values for participants with low, but not high synthesis capacity (*P* ≥ 0.021 for both). Error bars are within-subject SEM.

If dopamine mediates cognitive effort, it should be possible to increase motivation pharmacologically. Indeed, both methylphenidate and sulpiride increased subjective values for participants with low, but not high dopamine synthesis capacity (Fig. 1D; Fig. S2B—C). A mixed-effects model revealed that both methylphenidate (*β* = −0.069, *P* = 0.0042) and sulpiride (*β* = −0.10, *P* = 8.3×10^−4^) interacted with dopamine synthesis capacity to increase subjective values. Neither drug showed main effects (both *P* ≥ 0.37).

The converging effects of synthesis capacity and two separate drugs strongly implicate striatal dopamine. Methylphenidate blocks reuptake, increasing extracellular striatal dopamine tone (*23*) and can amplify transient dopamine signals (*24*). Sulpiride is a D2 receptor antagonist which, at low doses can increase striatal dopamine release by binding to pre-synaptic auto-receptors, enhancing striatal reward signals, and learning (*6, 25*). While sulpiride can block postsynaptic D2 receptors at higher doses (*26*), both drugs increased behavioral vigor (reaction times and saccade velocities; cf. (*6, 26*)), especially in low dopamine synthesis capacity participants, corroborating that both drugs increased dopamine release (Supplemental Results).

To assess whether dopamine amplified subjective benefits versus costs, we made a series of offers, in a second task, centered around participants’ indifference points (Fig. 2A). To generate specific predictions, we simulated psychometric choice functions with a computational model of striatal dopamine effects on decision making (*5*). With higher dopamine, the model predicts enhanced sensitivity to benefits and reduced sensitivity to costs. This manifests as a steeper choice function to the right of indifference, where the ratio of benefits to costs (of the high-versus low-effort option) is larger, but shallower functions to the left, where the benefits-to-costs ratio is smaller (Fig. 2B).

**Fig 2.**
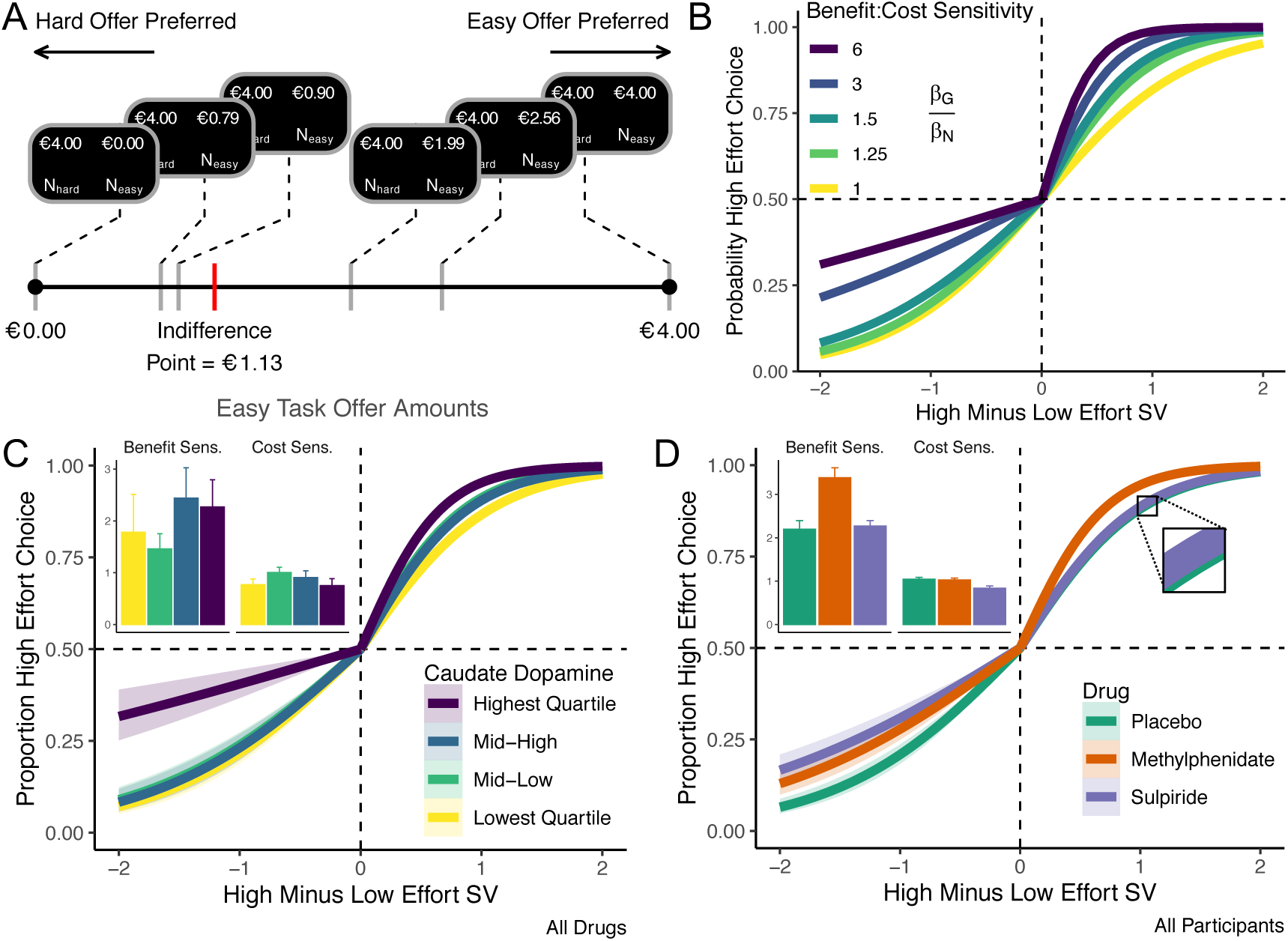
Dopamine alters valuation by re-weighting benefits-versus-costs of cognitive work. **A.** Low-effort (N_easy_) offers were paired with a €4 offer for a high-effort N-back task (N_hard_). **B.** Simulated effects of dopamine on benefit-versus-cost sensitivity are mirrored by empirical effects of **C.** dopamine synthesis capacity and **D.** pharmacological agents. Mixed-effects logistic regression curves and 95% CI fit across all drugs for each synthesis capacity quartile (**C.**) or all participants for each drug (**D.**). Insets show estimated effect of benefits and costs on choice across participants in (**C.**) each quartile +/- SEM and on (**D.**) each drug +/- within-subject SEM.

Choice behavior supported model predictions. Simulated effects were mirrored by effects of variability in dopamine synthesis capacity (Fig. 2C), and of methylphenidate and sulpiride versus placebo (Fig. 2D). Formally, high effort selection was sensitive to both benefits (offer amount differences; *β* = 2.30, *P* = 1.2×10^−9^) and costs (load differences; *β* = −1.07, *P* = 2.2×10^−16^). Critically, the effect of benefits increased with synthesis capacity (*β* = 0.65, *P* = 0.0024) and on methylphenidate (*β* = 1.34, *P* = 0.0048), while the effect of costs was attenuated on sulpiride (*β* = 0.24, *P* = 0.036). Participants also selected high-effort choices more often with higher dopamine synthesis capacity (*β* = 1.02, *P* = 3.1×10^−4^), and on methylphenidate (*β* = 1.75, *P* = 0.0016) versus placebo, but not reliably so for sulpiride (*β* = 0.46, *P* = 0.12). No other interactions or main effects were significant (all *P* ≥ 0.47).

These results clearly implicate dopamine in choice but they do not uncover how decision-making is altered. Dopamine could increase attention to benefits versus costs. Alternatively, it could alter the impact of these attributes on choice without impacting attention itself. We thus tracked eye gaze to quantify attention to attributes and how it interacted with dopamine. Proportion gaze at an offer (either costs or benefits) strongly predicted offer selection, (Fig. 3B; *β* = 0.30, *P* = 7.6×10^−6^; cf. (*27, 28*)). However, gaze at benefits predicted steeper increases in hard-task selection than gaze at costs (gaze by dimension interaction: *β* = 0.41, *P* = 1.1×10^−5^).

**Fig. 3.**
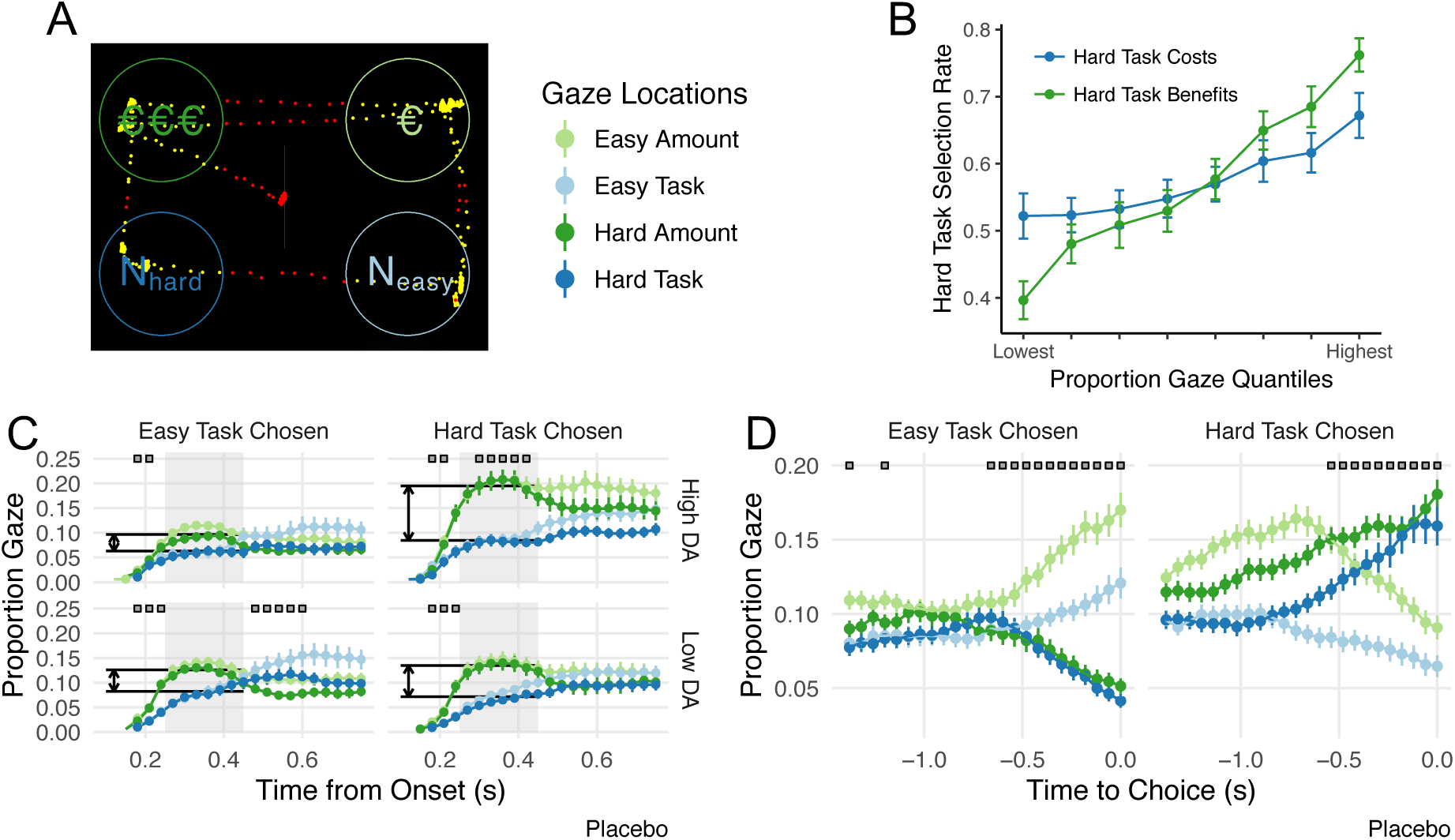
Effect of gaze, value, and dopamine synthesis capacity on effort selection. **A.** Participants decided between offers with costs (N-back load) and benefits (Euros) separated in space. Dots indicate gaze at (yellow) and away from (red) offers. **B.** Gaze predicted high-effort selection, and more so with gaze at benefits-versus-costs. **C—D.** Proportional (cross-trial) gaze at the four information quadrants following offer onset and leading up to response. **C.** Early gaze (250— 450ms following offer onset) indicated by grey shading. Boxes indicate time points at which participants gazed reliably more at either benefits or cost information (paired t-tests P < 0.05). **D.** Boxes indicate time points at which participants gazed reliably more at the selected offer (one-tailed paired t-tests, P < 0.05). All error bars reflect +/- SEM.

Gaze patterns implicated dopamine in enhancing the impact of attention to benefits-versus-costs on the decision to engage in cognitive effort. Early in a trial, participants fixated benefits (of either offer) more than costs and this asymmetry was larger on trials in which they chose the high-effort option (choice effect: *β* = 0.41, *P* = 0.0017; Fig. 3C). Moreover, this effect was stronger in participants with higher dopamine synthesis capacity (choice by synthesis capacity interaction: *β* = 0.37, *P* = 0.0045; top versus bottom row, Fig. 3C). For those with lower synthesis capacity, methylphenidate strengthened this relationship (interaction between drug, synthesis capacity, and choice: *β* = −0.36, *P* = 0.012; though sulpiride did not: *β* = −0.041, *P* = 0.78). Drugs and synthesis capacity did not impact gaze patterns themselves (*P* ≥ 0.10 for main effects), indicating that dopamine did not alter attention to benefits, but rather strengthened the impact of attention to benefits-versus-costs on choice.

Gaze may correlate with choice because attention amplifies the perceived value of attended offers, causally biasing choice (*27*). Alternatively, reversing this causality, participants may simply look more at offers they have already implicitly chosen (*28*). We found evidence for both: Early in a trial, attention influenced choice while, later, choice influenced attention. To address this, we fit drift diffusion models (*29*) in which cost and benefit information accumulate in a decision variable rising to a threshold. This variable is the instantaneous difference in the perceived value of the high-versus the low-valued offer. We considered ‘attention-biasing-choice’ models with multiplicative effects (i.e., gaze multiplies the effects of value information), and ‘choice-biasing-attention’ models in which gaze has a simple, additive effect (gaze correlates with choice but does not amplify value) (*28*). The best fitting model (Fig. 4A—C; Eqn. 1, *30*) included both additive and multiplicative effects (Supplemental Results).

**Fig. 4.**
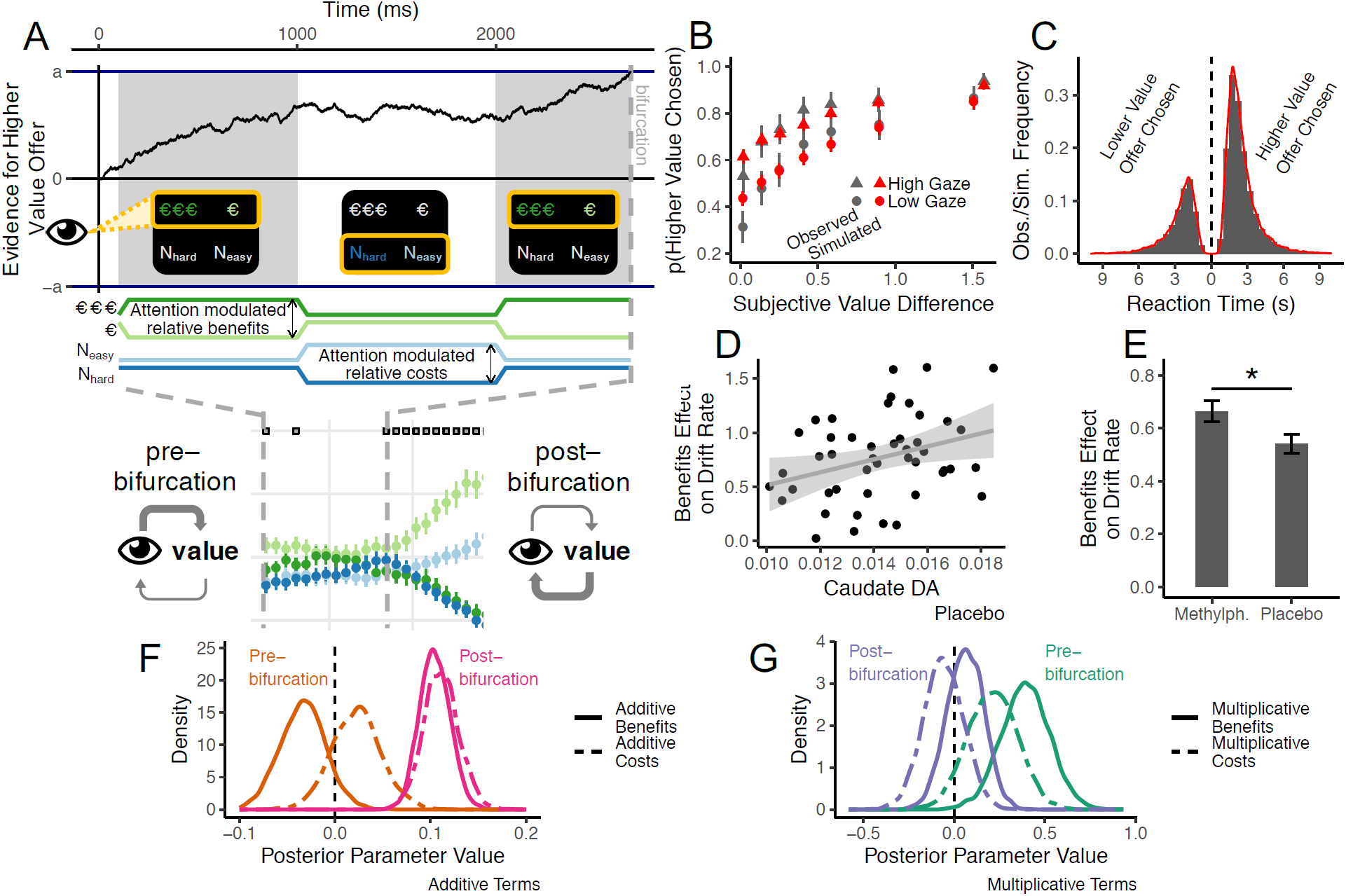
A. Gaze-attribute model: early gaze amplified the effect of attended-versus-unattended attributes on choice. Late gaze reflected the to-be-selected response. **B.** Model simulations (red) predicted choice (grey), and reaction time distributions (**C**). **D.** Benefits effect on drift rate correlate with dopamine synthesis capacity (95% CI shown). **E.** Methylphenidate enhances benefit effect. **F—G.** Posterior parameter densities from models fit alternately with pre- or post-bifurcation gaze on placebo. **F.** Additive benefit (*β*_1_ = −0.030; *P* = 0.076) and cost (*β*_2_ = 0.020; *P* = 0.81) gaze terms were approximately zero pre-bifurcation, and reliably positive post-bifurcation (*β*_1_ = 0.10; *P* < 2.2×10^−16^ and *β*_2_ = 0.11; *P* = 0.0031). **G.** Multiplicative interaction terms reveal that the effects of benefits (*β*_3_ − *β*_5_ = 0.40; *P* = 0.0024) and costs (at trend-level; *β*_4_ − *β*_6_ = 0.12; *P* = 0.060) were larger when fixating the respective attribute pre-bifurcation, while neither term was different from zero, post-bifurcation (*β*_3_ − *β*_5_ = 0.07; P = 0.27 and *β*_4_ − *β*_6_ = −0.060; *P* = 0.70). All error bars are +/- SEM.

We next considered the possibility that the gaze-value interactions changed dynamically across the trial. Indeed, ∼775 ms prior to responding participants began committing their gaze towards the to-be-chosen offer (Fig. 3D). Thus, while early gaze appeared to influence choice formation (Fig. 3C), later gaze appears reflect to latent choices, once formed. Based on this, we asked whether early attention causally amplifies attended attributes (a multiplicative combination), while late gaze simply correlates with choice (additive; Fig. 4A). To test our hypothesis, we split trials according to when participants began committing their gaze to the to-be-chosen offer (the “bifurcation”) for each participant and session and refit our model to gaze data from before or after this time point. The result supported our hypothesis. Multiplicative terms were reliably positive pre-bifurcation but near zero post-bifurcation, with the opposite pattern for additive terms (Fig. 4F—G). These results support that while early attention appeared to amplify the effect of benefits-versus-costs, later gaze simply reflected a latent choice.

Finally, we tested whether the effects of dopamine on choice could be attributed to these dynamic decision processes. Indeed, both higher dopamine synthesis capacity (on placebo; Eqn. 1: *β*_3_ + *β*_5_/2; Pearson *r* = 0.30, *P* = 0.039; Fig. 4D) and methylphenidate (*t*_*paired*_ 45 = 2.54, *P* = 0.015; Fig. 4E) increased the effect of benefits on evidence accumulation. The corresponding effect of sulpiride on cost was not significant (Eqn. 1: *β*_4_ + *β*_6_/2; *t*_*paired*_ 45 = −1.41; *P* = 0.17; Supplemental Discussion). We further found that methylphenidate amplified the effects of benefits on drift rate even when only modeling pre-bifurcation gaze (*t*_*paired*_ 45 = 2.44; *P* = 0.019) – prior to the latent choice. Collectively, our results support that striatal dopamine enhances motivation for cognitive effort by amplifying the effects of benefits versus costs attended early in a decision.

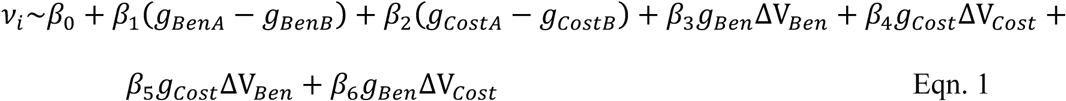

## Acknowledgments

We wish to thank the individuals who participated in this study, and James Wilmott for eye-tracking code and consultation.

## Author contributions

conceptualization: R.C., A.W., and M.J.F., data curation: J.I.M., R.v.d.B., L.H., D.P., formal analysis: A.W., R.v.d.B., R.C., M.J.F., funding acquisition: R.C., A.W., investigation: J.I.M., R.v.d.B., L.H., D.P., project administration: J.I.M. and R.C., software: A.W., writing: A.W., M.J.F, R.C., supervision: R.C. and M.J.F.;

## Data and materials availability

Data and analysis scripts will be publicly available at the conclusion of the parent study via http://hdl.handle.net/11633/aac2qvfx.

## Supplemental Methods and Results

### Supplemental Methods

#### Participants

50 Healthy, young adult participants (ages 18—43, 25 men) were recruited from The Netherlands to participate in a larger, within-subject, double-blind, placebo-controlled, pharmaco-imaging study. Participants were screened to ensure that they are right handed, Dutch-native speakers, healthy, and neurologically normal with no (relevant) history of mental illness or substance abuse. Participants were also excluded for history of hepatic, cardiac or respiratory disorders, epilepsy, hypersensitivities to methylphenidate, entacapone, carbidopa, or sulpiride, suicidality, smoking, diabetes, or claustrophobia. Participants also had normal or corrected-to-normal hearing and vision. Pregnant or breast-feeding participants were excluded. The study was approved by the regional research ethics committee (Commisssie Mensgebonden Onderzoek, region Arnhem-Nijmegen; 2016/2646; ABR: NL57538.091.16).

All complete datasets were included for analyses, as well as partial datasets, where available. PET data were not collected for two participants who were thus excluded from individual difference analyses testing the effects of dopamine synthesis capacity. These same two participants also failed to participate in all drug sessions and were excluded from analyses of relevant drug effects. One did not participate in either the placebo or methylphenidate session, while the other did not participate in the methylphenidate session. In addition, a third participant did not participate in the sulpiride session. While remaining participants completed all sessions, two more participants showed no sensitivity to cognitive demands in the discounting task, never once selecting the low-effort, low-reward option in any drug session. Given uncertainty about whether these participants followed task instructions to consider both choice dimensions, these two participants were excluded from all analyses.

#### General Procedure and Tasks

The broader study (n = 100 participants) was designed to investigate the effects of dopaminergic drugs on cognitive control, and how those drug effects depend on baseline dopamine synthesis capacity. Participant engagement spanned five visits: a 3-hour screening session, three, 6-hour pharmaco-imaging sessions with multiple tasks both in, and out of an fMRI scanner after being administered placebo, sulpiride, or methylphenidate, and a final 2.5-hour PET session for measuring dopamine synthesis capacity. Errors in drug scheduling meant that drug session order was not perfectly counterbalanced. Consequently, 23, 15, and 10 participants took placebo on session number 1, 2, and 3, respectively, while the numbers were 12, 18, and 18 for sulpiride, and 13, 15, and 20 for methylphenidate. Given data loss and imperfect counterbalancing of drug by session order, we confirmed all inferences via hierarchical regression analyses controlling for session order as a factor.

During screening, after providing written consent, participants completed medical and psychiatric screening interviews, reviewing height, weight, pulse rate, blood pressure, and electrocardiography, neuropsychological status, and existence of (relevant) DSM-IV axis-I disorders, and ADHD. Next, participants completed a structural T1-weighted magnetization prepared, rapid-acquisition gradient echo sequence MRI scan (TR 2300 ms, TE 3.03 ms, flip angle 8°, 192 sagittal slices, 1 mm thick, field of view 256 mm, voxel size 1×1×1 mm), scanned by a Siemens MAGNETOM Skyra 3 Tesla MR scanner. Finally, participants completed digit span and a listening span working memory tests, and we also measured their resting eye-blink rate via electrooculography.

Participants were asked to refrain from smoking or drinking stimulant-containing beverages the day before a pharmaco-imaging session, and from using psychotropic medication and recreational drugs 72 hours before each session, and were also required to abstain from cannabis throughout the course of the experiment. During a pharmaco-imaging session, participants completed multiple tasks, including the tasks which are the focus of this study: the N-back task, a cognitive effort discounting task adapted from (*2*), and a gaze-decision making task. At the beginning of a session, participants completed another screening form and a pregnancy test. In addition, we measured baseline subjective measures, mood and affect, as well as temperature, heart rate, and blood pressure at baseline (these measures were also recorded at two fixed time points after drug administration). We further monitored baseline mood and affect before and after drug administration. Other tasks which participants completed, but which were not analyzed here, included a reinforcement learning task designed to dissociate contributions of reinforcement learning and working memory during stimulus-response learning, and three tasks measuring creativity. Participants also completed two tasks in the fMRI scanner: one measuring striatal responsivity to reward cues and a reversal learning task. Finally, we also collected measures of depression, state affect, BIS/BAS, impulsivity, and the degree to which participants pursue cognitively demanding activities in their daily life.

At least two days after the pharmaco-imaging sessions, during the PET session, participants also performed a Pavlovian-to-instrumental transfer task and collected a general intelligence measure (a WAIS IV fluid intelligence test). A preregistration for the broader study, as well as a complete list of measures collected, and their intended use is detailed in a pre-registration: https://www.trialregister.nl/trial/5959.

Note that the first 50 participants recruited for the broader (n = 100) study also completed a different, yet complementary decision-making task in which participants decided whether to engage with a demanding, but rewarded working memory task, or instead have free time (Hofmans, Papadopetraki, van den Bosch, Määttä, Froböse, Zandbelt, Westbrook, Verkes, & Cools, 2019, bioRxiv).

All tasks analyzed in this paper were presented using Psychtoolbox-3 for MATLAB. Prior to drug administration, participants completed all levels of the N-back task to re-familiarize themselves with the subjective demands of each level. The N-back task was performed off-drug so that drugs would alter neither performance nor subjective experience of the N-back. Participants were administered drugs prior to the effort discounting and gaze-decision making tasks. To accomplish double-dummy blinding as to drug condition, participants took one capsule at each of two different time points: time point one was either placebo or 400 mg sulpiride, while time point two was either placebo or 20 mg methylphenidate. 50 minutes after taking methylphenidate (or placebo on sulpiride and placebo days), 140 minutes after sulpiride (or placebo on methylphenidate and placebo days), or 50 after taking the second placebo (on placebo days), participants performed the effort discounting and gaze-decision tasks. These times were chosen to maximize the impact of drugs which near their pharmacokinetic and physiological effect peaks in the range of 60—90 and 60-180 minutes for methylphenidate (*31*) and sulpiride (*32*), respectively.

#### N-back Task

For the N-back task, off-drug, participants completed levels N = 1—4, performing three rounds of each level. Each round comprised a series of 40 upper-case consonants presented for 2.5 seconds, during which participants were required to respond by button press indicating whether each letter was a “target” or “non-target”. After response, the stimulus was replaced by a central fixation cross until the subsequent stimulus was presented, 3 seconds after the last stimulus onset. Each N-back level was referred to by one of four lower-case vowels (‘the *a* task’ for the 1-back, ‘the *e* task’ for the 2-back, etc.). Vowels were used as task labels rather than explicit, numeric representations for each level to avoid anchoring confounds while participants considered (numeric) subjective values in the subsequent discounting and gaze-decision trials.

#### Discounting Task

During the discounting task, participants were asked, on-drug/placebo, to choose between repeating one of each of the higher levels of the N-back task (N = 2—4) for one of two larger amounts of money (€2 or €4) and a lower level (N = 1—2) for a smaller, variable amount of money on each trial. On the first trial for each high-effort / low-effort pair the initial offer for the low-effort offer was one half the offer for the high-effort offer. After each choice, the offer amount for the low-effort offer was adjusted down if it was chosen, and adjusted up if it was not chosen until the participant was indifferent between the offers. The magnitude of the adjustment was half as much on each trial such that the offer converged to the indifference point over 5 trials (i.e., until offers are within €0.0625 of the assumed indifference for €4 offers and within €0.03125 for €2). The adjusted offer after the fifth decision trial was taken to be the indifference point, and this quantified the subjective value of the high-effort, relative to the low-effort offer. For example, if a participant were indifferent between €4 for the 3-back, and €1.13 for the 1-back, the subjective cost of the 3-versus the 1-back is €2.87, and the subjective value of the €2 offer for the 3-back was €1.13. Note that when testing the effect of the offer amount on subjective values, we normalized the subjective value by the base offer (e.g. €1.13 / €4 = 0.2825). In total, participants completed 50 discounting trials comprising 5 decision trials for each of 5 high-/low-effort pairs and each of 2 base offer amounts. All discounting trials were self-paced.

#### Gaze-Decision Task

After we established indifference points, participants completed an additional 168 self-paced choice trials while we monitored their gaze. Offers were tailored to participants’ indifference points to alternately bias high-cost / high-benefit offer selection, or low-cost / low-benefit selection on half of the trials. Offers were further designed to manipulate choice difficulty, with trials varying from difficult discriminations – in which offers were close in subjective value, to easy discrimination trials – in which subjective offer values were maximally different. This design ensured that we sampled from across the psychometric choice function, but also emphasized difficult discrimination trials maximizing sensitivity to, for example, subtle drug and gaze effects on choice. Specifically, we included 18 easy discrimination trials to ensure that participants were paying attention: 9 in which we offered either the same offer amount for the easy and hard task (participants should mostly choose the easy task), and 9 in which we offered €0 for the low-effort offer (participants should mostly choose the hard task). Indeed, as anticipated, participants overwhelmingly selected the higher value offer on these easy decision trials, whether that offer was the high-effort alternative (94.2% of trials across all drugs and participants) or the low-effort alternative (90.1% of trials). We also included 150 difficult discrimination trials: 75 in which we offered 20—30% below the indifference point for the low-effort offer (percentage sampled from a uniform distribution spanning the range), and another 75 trials in which we offered 30—50% of the difference between the indifference point and the high-effort offer above the indifference point (see Fig. 2A for an example set of offer pairs). These ranges were used because prior piloting revealed that they were close enough to indifference that participants made choices contrary to offer biases at a desired rate (we designed our ranges to achieve a 20—30% rate of “anti-bias” trials; overall participants chose against offer biases 30.6% of the time). The ranges for bias high-effort and bias low-effort trials were asymmetric because prior piloting further revealed that for a given range (% difference from the indifference point), participants tended to select the high-effort offers at a higher rate, on trials in which we biased high-effort selection, than the rate at which they selected the low-effort offers on trials in which we biased low-effort selection. Indeed, this is expected if participants are relatively more sensitive to benefits than costs: for a given range, participants will tend to select the high-effort option more often on bias high-effort trials because the psychometric choice function has a steeper slope than on bias low-effort trials (see Fig. 2). Thus, we increased the range to more strongly bias low-effort selection on bias low-effort trials to achieve greater balance in overall rate of high- and low-effort selection rates. Participants’ choices reliably reflected offer biases. However, the final anti-bias choice rate on difficult, bias high-effort trials was 18.9%, and on bias low-effort trials was 42.4%, indicating that the propensity to select high-effort offer was not fully offset by the stronger percentage range of offer biases used to bias low-effort selection.

Participants had up to 9 seconds to indicate their preference by button press, after offer onset, before a trial would time-out and advance to the next trial. Across all sessions and participants, only 0.059% of trials (6 trials on placebo, 4 trials on methylphenidate, and 8 on sulpiride; 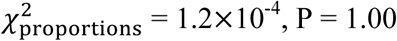) timed-out and were excluded from analyses. When participants responded, the selected offer was highlighted by a rectangular frame for 0.5 seconds, and then offers were replaced by a central fixation cross indicating the start of the next trial. Participants’ decision trials were broken up into 3 runs of 56 trials each with breaks for rest and to recalibrate eye tracking in between.

#### Eye Tracking

Participants’ gaze and saccade vigor was monitored during the gaze-decision task using an Eyelink 1000 infrared camera (SR Research; Ottawa, Ontario). Participants rested their head on a table-mounted chin rest with their eyes approximately 76 cm from a 61 cm LCD monitor; gaze position readings were recalibrated at the beginning of each run of decision trials. At the beginning of each choice trial, a central fixation cross was presented, on which participants were required to fixate for 1 second to initiate the trial. After successfully holding fixation for 1 second, two offers were presented, each comprising an amount in Euros, and an N-back task level, with the four pieces of information displayed in the four corners of the screen. Offers were left-right lateralized, while cost (e.g. ‘*a*’, ‘*e*’, etc.) and benefit (e.g. €1.70 and €2.00) information was presented on either the top or bottom on each trial. Positions of the costs versus benefits and side of the high-cost, high-benefit offer were selected randomly on each trial. Each piece of information was centered 11 degrees away from the central fixation cross, and subtended between approximately 0.37—1.37 degrees of visual angle. Gaze position was sampled every 0.003 seconds, and was down-sampled to every 0.01 seconds for analyses.

To identify fixations, we considered both the sustained duration and location of gaze samples. First, samples within approximately 23 degrees of visual angle of the stimulus centroid (fully encompasses “central” and “near peripheral” vision) were tagged as directed at the relevant choice feature. Then, we counted any interval of gaze directed continuously at the same feature for longer than 70 ms as a fixation, reasoning that anything shorter would be well below minimum duration of typical fixations and must reflect a passing saccade. These liberal thresholds ensured that we counted every possible sample of gaze that may have contributed to information acquisition towards our proportion gaze measures. Fig. 3A shows a typical example trial and gaze samples counted as either at, or away from offer information.

To compute saccade vigor, we calculated the degrees of visual angle between each successive time point (from the down-sampled 100 Hz samples). Next, we identified saccades as sequences of successive time points differing by more than 100 degrees of visual angle per second. Next, we identified the peak velocity in each of these sequences. Finally, we regressed out the saccadic main sequence by taking the residuals of a linear model regressing peak saccade velocities onto saccade distances across all saccades. We used these residualized peak velocities in our analyses of the effects of striatal dopamine on saccadic vigor.

#### PET Imaging

We used the radiotracer [^18^F]-fluoro-DOPA (F-DOPA) and a Siemens mCT PET-CT scanner to measure participants’ dopamine synthesis capacity. Images were captured using 40 slice CT, 4 x 4 mm voxels, with 5 mm slice thickness. One hour prior to F-DOPA injection, participants received 150 mg carbidopa to reduce decarboxylase activity and 400 mg entacapone to reduce peripheral COMT activity with the intention of increasing the bioavailability of the radiolabeled F-DOPA and enhance signal to noise. Following the Pavlovian-to-instrumental task, then entacapone and carbidopa administration, participants performed a cognitive task battery while waiting for peak drug efficacy. About 50 minutes after administration, participants were positioned to lie down comfortably and a nuclear medicine technician administered a low dose CT to correct attenuation of PET images. Subsequently, participants were administered a bolus injection of 185 MBq (5 mCi) max F-DOPA into the antecubital vein. Over the course of 89 minutes, we then collected 4 1-minute frames, 3 2-minute frames, 3 3-minute frames, and 14 5-minute frames. Data were reconstructed with weighted attenuation correction, time-of-flight correction, correction for scatter, and smoothed with a 3 mm full-width-half-max kernel.

Data were preprocessed using SPM12. All frames were realigned to the middle (11th) frame to correct for head movement. Realigned frames were then co-registered to the structural MRI scan, using the mean PET image of the first 11 frames, which have better contrast outside the striatum than the later frames. Presynaptic dopamine synthesis capacity was quantified as F-DOPA influx rate (Ki; min^-1^) per voxel using Gjedde-Patlak linear graphical analysis (*33*) for the frames of 24—89 minutes. These Ki values represent the amount of tracer accumulated relative to the reference region of cerebellum grey matter. The reference region was obtained using FreeSurfer segmentation of each individual’s high resolution anatomical MRI scan. Ki maps were spatially normalized to MNI space and smoothed using an 8 mm FWHM Gaussian kernel.

After preprocessing and normalization to MNI space, Ki values were extracted from masks defining regions of interest based on an independent, functional connectivity-based parcellation of the striatum (*20*). In particular, we extracted Ki values from 3 striatal regions – the caudate nucleus (817 voxels), the putamen (1495 voxels), and the ventral striatum / nucleus accumbens (607 voxels), and averaged across all voxels in each region for individual difference analyses. Our individual difference analyses focus on the caudate nucleus. All results survive Bonferroni correction across the three striatal sub-regions with the sole exceptions being the impact of caudate nucleus Ki on the benefits effect on the drift rate (P × 3 = P_Bonferroni_ = 0.12) and the effect of Ki on subjective values in the placebo session (P_Bonferroni_ = 0.066; along with lower-power AUC analyses collapsing across load levels and offer amounts). Nevertheless, the influence of caudate Ki on subjective values in the discounting phase is confirmed by hierarchical, trial-wise regression analyses across sessions (P_Bonferroni_ = 0.022), revealing robust effects, surviving correction for multiple comparisons.

#### Simulating Dopamine’s Effects on Sensitivity to Costs and Benefits

As noted in the main text, we tested the hypothesis that striatal dopamine has asymmetric effects on benefits versus costs sensitivity during decision-making. To simulate these effects, we adopted the Opponent Actor Learning Model (OpAL; *5*) according to which a subjective action value is given by a linear combination of costs and benefits, where the cost and the benefits terms have distinct weights (*β*_*N*_ and *β*_*G*_, respectively). To model our decision-making task, we thus consider the subjective value of an offer (*V*_*p*_) during the gaze-decision task to be:

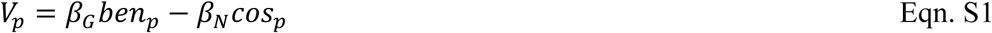

Here *ben*_*p*_ is the benefit of offer *p* in terms of objective monetary amount (€), and *cos*_*p*_ is the objective N-back task level. The weights thus convert objective measures into subjective benefits and costs and can moreover be modulated independently (e.g., by dopamine). Following (*5*), we simulated increases in dopamine release as an increase in the ratio of *β*_*G*_ to *β*_*N*_ (Fig. 2A). We then mapped values to choice probabilities via softmax, such that the probability of choosing the low-effort offer (*lo*) versus the high-effort offer (*hi*), is:

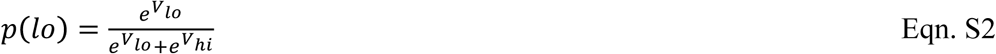

To specifically simulate the choice probability functions in Fig. 2B, we assumed a high-effort offer amount, and a fixed difference in costs, and computed the low-effort offer amount required for indifference for a given ratio of *β*_*G*_ to *β*_*N*_. Next, we computed the low effort offer amount required for a given proportional shift along the x-axis (the difference in subjective values), as the fraction of the distance between the indifference point and the low-effort offer bounds: €0—*ben*_*hi*_. Finally, we computed the subjective value of this low-effort offer using Eqn. S1 and the probability of the high-effort offer selection (*p*(*hi*) = 1 − *p*(*lo*)) using Eqn. S2.

#### Hierarchical Regression Analyses

All hierarchical regression models were fully random and fit using the lme4 package version 1.1-17 for R. The following were reported in the main text or Supplemental Results.

For the discounting task, we estimated the effect of z-scored high-effort offer amount (*amt*), drug as a factor (*drug*), z-scored load difference (*ldiff*), and session number as a factor (*S*; in the following equation, session is dummy coded depending on which session the trial comes from, e.g. if Session 2: *S*_2*i*_ = 1, *S*_3*i*_ = 0, etc.) on the subjective value (*SV*_*ij*_; given here as the indifference point divided by the offer amount) of a high-effort offer *i*, for participant *j* by fitting the following hierarchical regression model:

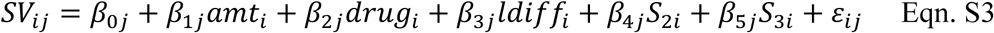

Additionally, while all terms have subject-specific intercepts, we also allowed slopes to vary by participant, with z-scored caudate dopamine synthesis capacity (*cDA*) as a subject-level predictor of the intercept and drug terms, thus modeling a cross-level interaction of dopamine synthesis capacity and drug status:

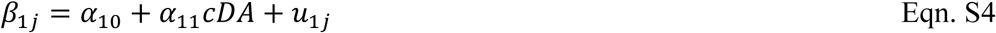

and a main effect of dopamine synthesis on subjective value.

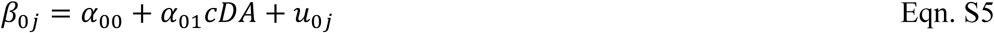

Note that *u* and *ε* are error terms. A full list of fitted model fixed effect parameters and standard errors is provided in Table S1.

For the gaze-decision task, we fit a hierarchical logistic regression to estimate the effects of z-scored relative (high-effort offer versus low-effort offer) benefits (*ben*) and costs (*cost*), caudate dopamine synthesis capacity, and drug status on binary choice (*Chc*) of the high-effort offer on trial *i* for participant *j*, controlling for session number.

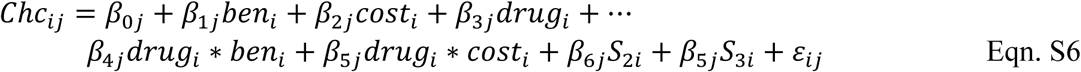

Note that we specified our intercept term in the same way as in the previous hierarchical regression, allowing for a cross-level main effect of dopamine synthesis capacity (Eqn. S5). In addition, we allowed for cross-level interactions to test how dopamine synthesis capacity modulated both benefit and cost terms, *k*. Fitted model results are provided in Table S2.

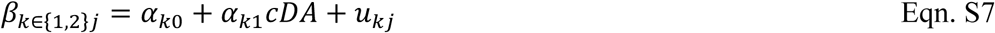

We also modeled the effects of costs and benefits as a function of dopamine synthesis capacity quartiles or as a function of drug in separate models, to visualize the relevant interactions (Fig. 2B—C, insets). For modeling the effect of dopamine synthesis capacity as quartiles, we fit a simplified version of Eqn. S6, in which we modeled separate benefit and cost terms for each dopamine synthesis capacity quartile, *q*.

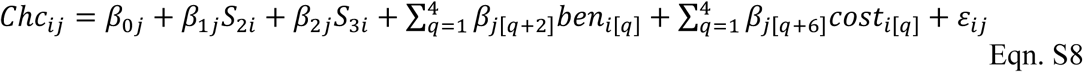

Similarly, we modeled benefit and cost effects on each the dopamine drugs versus placebo (methylphenidate, MPH, and sulpiride, SUL).

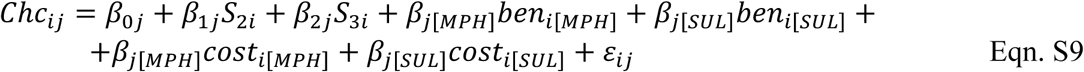

Next, we computed participant-specific effect estimates by summing the corresponding random effects, and fixed effects (for their respective quartile, e.g. *β*_*j*[*q*+2]_, or for the respective drug, e.g. *β*_*j*[*MPH*]_). Finally, we plotted the mean and standard error of these estimated effects in the insets of Figs. 2B—C. We also plotted individual participants’ benefit and cost effect estimates in the Supplemental Results (Fig. S5).

With respect to gaze in the gaze-decision task, we first fit a hierarchical logistic regression to understand how gaze and offer values influenced choice. Specifically, we tested for effects of z-scored proportion dwell times on the high-effort offer (*hG*: high-effort offer dwell time minus low-effort offer dwell time, normalized by total dwell time) and of z-scored combined offer value (*totSV*), and their interaction, as well as the difference in subjective values (*diffSV*: high-effort minus low-effort subjective value), and session number predicting binary choice (*Chc*). Fitted model results are provided in Table S3 and described in the Supplemental Results under “*Effects of Gaze and Attribute Values on Choice”*.

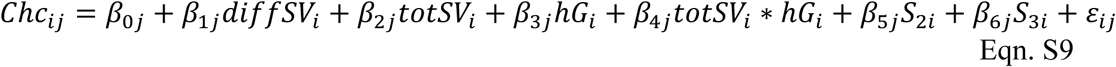

To analyze the relationship between dynamic gaze patterns, dopamine, and choice, we began by estimating average fixation patterns at every time point following offer onset (the frequency of fixating one of the four information quadrants) across trials, for every subject. We computed separate averages for each drug session to test for relationships with dopamine, and furthermore computed separate averages for choose-high versus choose-low effort trials so that we could test for a relationship with choice. These averages, when further plotted as means and standard errors across participants, revealed a clear pattern of preferential gaze at benefit versus cost information in an interval between 250 ms and 450 ms after offer onset (Fig. 3C). We then asked whether this pattern differed by dopamine status and choice by fitting a hierarchical regression testing whether benefits versus cost gaze averages (*bcG*_*i*_: proportion of trials looking at benefits versus cost information, averaged across all time points in the 250—450 ms window, for average *i*) varied by choice type (*cType*: high-effort versus low-effort chosen) and drug as within-participant factors and dopamine synthesis capacity as a between-participants continuous predictor. Here, as above, all first-level predictors vary by participant (*j*). Fitted model results are provided in Table S4.

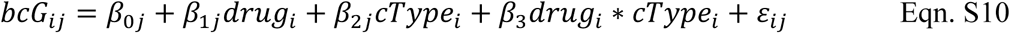

As above, we allowed for randomly-varying main effects of caudate dopamine synthesis capacity (Eqn. S3). We further allowed the effect of drug and choice type to vary randomly by dopamine synthesis capacity (as in Eqn. S5), and the drug by choice type interaction to vary non-randomly by caudate synthesis capacity.

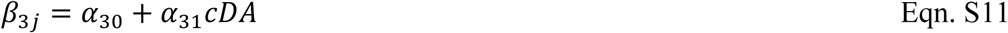

Given the number of averages we fit in this model (one for each subject, drug session, and choice type), allowing the drug by choice type interaction to also vary randomly by subject required too many degrees of freedom to be estimated. However, the negative three-way interaction we observed (dopamine synthesis capacity by choice type by methylphenidate; Table S4), is entirely consistent with complementary, non-hierarchical models we estimated separately for the methylphenidate and placebo sessions. In those models, we found that the two-way interaction between dopamine synthesis capacity and choice type predicting higher average gaze at benefits versus costs on methylphenidate was an order of magnitude smaller (*β* = 0.037; P = 0.86) than it was on placebo (*β* = 0.44; P = 0.031).

#### Drift Diffusion Modeling

To understand how value and gaze combine to influence evidence accumulation during choice, we used the Hierarchical Drift Diffusion Modeling (HDDM) package (*29*). HDDM utilizes Markov Chain Monte Carlo sampling for Bayesian estimation of both group- and participant-level parameters (drift rate, threshold, etc.). Since our primary questions were about how drift rate varied across trials, we used the HDDMRegressor method which enables specifying trial-wise predictors of DDM parameters, to ask how drift rate varied by gaze and value measures.

To adjudicate between competing models by which gaze either amplifies the effect of attended versus unattended values on choice, or merely reflects implicit preferences, we fit drift diffusion models in which we allowed the drift rate to vary, respectively, by either multiplicative or additive combinations of gaze and value. Moreover, we also sought to adjudicate between competing models by which choice is driven by visual attention to either alternative offers, or offer attributes. Thus, competing models had drift rate varying by interactions of gaze and either net offer values (benefits minus costs) or attribute values.

Note that all trials were modeled except for non-response trials and trials in which participants responded too rapidly – based on a cutoff of reaction times > 250 ms. Only 17 out of 23767 trials were thus excluded from the HDDM modeling, or 0.071%.

The first, simplest model we considered had an additive combination of proportional gaze (proportion of total gaze at any piece of information in a trial) at offers *A* (*g*_*A*_) versus *B* (*g*_*B*_) and net offer values (*V*_*A*_ and *V*_*B*_) predicting drift rates towards offer *A* for participant *j* on trial *i* (*ν*_*i*_). Note that *A* was the higher value offer (regardless of whether it was the high-cost, high-benefit, or low-cost, low-benefit offer).

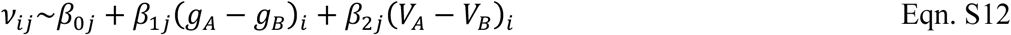

We also considered a model in which gaze at offers and net offer values interacted multiplicatively. This model is equivalent to the attention drift diffusion model (aDDM) of gaze-value interactions whereby gaze discounts the value of the unattended relative to the attended offer (*27*). In this model, the final term (*β*_2_) gives the effect of the value difference between *A* and *B* as a function of looking at *B* versus *A*, and, as noted in prior work (*28*), the ratio *β*_1_/*β*_2_ gives the fraction by which an offer is discounted when it is unattended relative to when it is attended.

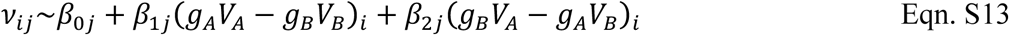

Next, we considered variants of these two models where visual attention is directed at offer attributes: gaze at the benefits (*g*_*BenA*_) and the costs (*g*_*CostA*_) of offer *A*. Here, the additive model takes the following form.

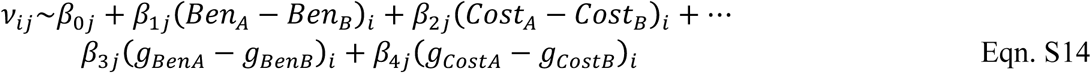

The interactive, attribute-wise variant model is given by the following.

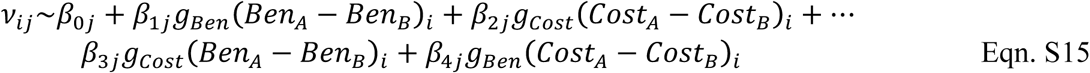

We furthermore considered a model which had both additive and multiplicative combinations of gaze and value. The net value model is identical to Eqn. S13, with the addition of a simple gaze term. This model captures the possibility that gaze and value combine dynamically across the trial (e.g. multiplicatively early in a trial as gaze amplifies value differences and additively late in a trial as gaze comes to reflect preferences as they form).

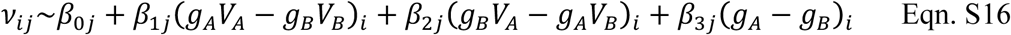

Finally, we also considered a model with both additive and multiplicative combinations of gaze and attribute values. This model, as noted in the main text, was the winning model with respect to AIC values across all sessions, and across individual drug sessions as well (Table S5).

AIC scores for data estimated across all sessions and in each session individually revealed that net value models consistently fit worse than attribute-wise value models, regardless of whether the models involved additive or multiplicative combinations of gaze and values. Also, while additive models consistently performed better than purely multiplicative models, the scores for combined additive plus multiplicative models were always best. Furthermore, supporting our hypothesis that gaze and value combine multiplicatively early in a trial, and additively late in a trial, we found that AIC scores were better for the multiplicative and multiplicative plus additive models based on pre-bifurcation gaze data and better for the additive model based on post-bifurcation gaze data (Table S6).

In addition to AIC scores, key qualitative features of our gaze data support a combined additive plus multiplicative model. First, the effect of gaze at costs on choice argues for either an additive or at least an additive plus multiplicative model rather than a purely multiplicative model. Namely, because load discounted the subjective value of offers, and the cost attribute necessarily carries a negative subjective valence, gaze at costs should discourage high-effort selection. A purely multiplicative model predicts that the more participants fixate a negative attribute, they less likely they should be to choose it. Nevertheless, we found clear evidence that the more participants fixated the costs of the high-cost, high-benefit offer, the more likely they were to choose it (Fig. 3B). Thus, a purely multiplicative model would not capture the effect of gaze at cost information. And yet, qualitative gaze and choice patterns also support multiplicative contributions. Specifically, for example, a fully-random, hierarchical regression of gaze and attribute values on choice reveals that the effect of costs on choice grows the more participants fixate costs. Fitted model weights show that selection of the higher-valued offer is predicted by the relative benefits of the offer (*β* = 4.4, P < 2.2×10^−16^), the relative costs (*β* = - 3.3, P < 2.2×10^−16^), and interactions reveal that while increasing proportion gaze at costs does not modulate the effect of benefits on choice (*β* = 0.14, P = 0.35), proportion gaze at costs did reliably increase the effect of costs on choice (*β* = −0.40, P = 0.0056). Thus, multiplicative combinations are needed to account for these types of interactions.

In addition to comparing quantitative and qualitative measures of model fit, we also performed posterior predictive checks to ensure that our selected model could reproduce our data. To do so, we used our selected model to simulate 500 data sets for every trial and confirmed that statistics of our observed data matched expectations from the simulations. For choices, we ensured that simulations closely matched the observed rate at which participants selected the higher-valued offer as a function of offer value difference and above-versus below-median proportion gaze at the higher valued offer (Fig. 4B). We also ensured that the following observed statistics matched our simulations for reaction times: the 10^th^, 30^th^, 50^th^, 70^th^ and 90^th^ percentile of the reaction time distributions, as well as the standard deviation of the mean reaction time, separately for distributions in which the participant did and did not select the higher-valued offer on each trial. Furthermore, a comparison of simulated and observed reaction time distributions for trials in which participants selected the higher and lower valued offers demonstrates excellent agreement between the model and the data (Fig. 4C).

#### Breaking up Trials According to Gaze Bifurcation

To test the hypothesis that gaze and value multiplicatively interact early in a trial and combine additively late in a trial, we broke trials into early and late gaze phases according to the time at which participants, on average, begin to commit their gaze to the chosen offer, prior to responding. We used a peak-finding method to identify the point at which participants’ gaze towards the unchosen option peaked, on average, for every subject and every drug session, before declining. For each participant and each session, we first computed timeseries averaging the proportion of trials fixating either the high- or low-effort offer, at every time point, time-locked to response. Next, we smoothed each of these timeseries using a two-sided linear filter. Then, we found the time point corresponding to the maximum proportion of trials fixating the unchosen offer in the 2 seconds prior to response. Across participants, the mean early-late split on placebo occurred, on average, 776 ms prior to response (SD = 360 ms), on methylphenidate it occurred 864 ms prior (SD = 344 ms), and on sulpiride it occurred 746 ms prior to response (SD = 306 ms). There was no difference in the mean split time between placebo and either methylphenidate (t_paired_ = 1.46, p = 0.15) or sulpiride (t_paired_ = −0.51, p = 0.61). However, bifurcation was earlier (with respect to the response time) on methylphenidate than on sulpiride (t_paired_ = 2.61, p = 0.011). We then split each participants’ gaze data according to whether samples were recorded before or after participant-specific split times prior to response. On trials with response times faster than participant-specific split times, we simply cut trials in half.

### Supplemental Results

#### Discounting by Drug and Caudate Nucleus Dopamine Synthesis Capacity

Participants with below-median dopamine synthesis capacity in the caudate nucleus discounted more steeply at every level of the N-back task in comparison with participants with above-median dopamine synthesis capacity, on placebo (Fig. S1A). This analysis is consistent with our area under the curve analysis reported in the main text. Also, there were no group differences on methylphenidate or sulpiride at any level as both drugs reliably increased the subjective offer values / indifference points selectively for participants with low dopamine synthesis capacity (Fig 1D), thereby erasing group differences (Fig. S1B—C).

#### Discounting by Dopamine Synthesis Capacity Outside the Caudate Nucleus

Outside of the caudate nucleus, dopamine synthesis capacity (Ki) in neither the putamen nor the ventral striatum / nucleus accumbens showed robust relationships with subjective offer values (Fig. S2). As reported in the main text, during the discounting phase, caudate nucleus Ki predicted subjective values (controlling for session order, Eqn. S3; *β* = 0.070, P = 0.0072) and these individual differences interacted with both methylphenidate (*β* = −0.069, P = 0.0042) and sulpiride versus placebo (*β* = −0.10, P = 8.3×10^−4^). By contrast in the putamen, there were no main effects of Ki (*β* = 0.037, P = 0.17), and no reliable interactions with either methylphenidate (*β* = −0.042, P = 0.10) or sulpiride versus placebo (*β* = −0.054, P = 0.096). Finally, in the nucleus accumbens, there was no main effect of Ki (*β* = 0.011, P = 0.69), and no reliable interaction with sulpiride (*β* = −0.034, P = 0.27). There was a negative interaction between Ki and methylphenidate versus placebo (*β* = −0.059, P = 0.029) in the nucleus accumbens, however this result does not survive Bonferroni correction. Voxel-wise analyses are presented (Fig. S1) to show main effects of Ki on placebo and interactions with methylphenidate and sulpiride versus placebo in all areas with high F-DOPA uptake signal.

We were open to the possibility that Ki in other regions predict willingness to accept offers to perform high-cognitive effort tasks for money. However, the caudate nucleus was a particularly strong candidate. First, the caudate nucleus has traditionally been regarded as “cognitive” (as opposed to “motor”) striatum and indeed fMRI work supports that cognitive motivation is encoded specifically in the caudate nucleus, while physical motivation was encoded more specifically in the putamen (*7, 21*). Second, a large-scale coactivation analysis of 5,809 fMRI datasets found that the anterior caudate nucleus was most commonly implicated in incentivized behavior, while the posterior caudate was most commonly implicated in executive function (*22*). Thus, the literature tends to support the hypothesis that the caudate nucleus plays a particular role in incentivized cognitive control. By contrast, the putamen was most commonly implicated in sensorimotor processes and the social and language-related functions (*22*). Also by contrast, the human ventral striatum was implicated in more non-specific forms of incentive and value processing by the fMRI coactivation meta-analysis (*22*). While our results do not implicate ventral striatal dopamine in promoting cognitive effort, they do not strictly rule out its involvement either.

#### N-back Performance

The N-back task features parametrically increasing working memory demands. As anticipated, we found that performance declined monotonically with increasing load, using the sensitivity index d’ (Table S7). The d’ measures performance while controlling for a response bias (in the N-back, d’ is a standardized measure of hit minus false alarm rate for N-back repeats). According to a mixed effects regression, d’ decreased linearly with load (*β* = - 0.69, P < 2.0×10^−16^), but, as anticipated, does not vary by drug session (methylphenidate: *β* = 0.053, P = 0.72; sulpiride: *β* = −0.16, P = 0.39), and there is no load by drug interaction (methylphenidate: *β* = 0.015, P = 0.76; sulpiride: *β* = 0.089, P = 0.14). Given that drugs were taken after N-back performance, no drug effects on N-back performance were expected.

We were open to the possibility that declining N-back performance could contribute to task aversion and thus discounting. However, we were also interested in whether dopamine status would explain differences in discounting, controlling for performance. To address this question in our own data, we fit the same mixed effects model from the main text (Eqn. S3), testing whether subjective values could be predicted by load, offer amount, dopamine synthesis capacity, and drug status, and added d’ as a covariate to control for performance. Critically, while higher performance predicted higher subjective values (*β* = 0.12, P = 7.0×10^−8^), we found that individual differences in dopamine synthesis capacity remained a positive predictor at trend level (*β* = 0.043, P = 0.082) and moreover both methylphenidate (*β* = −0.053, P = 0.038) and sulpiride (*β* = −0.088, P = 0.0096) still reliably interacted with dopamine synthesis capacity. These results indicate that dopamine-related changes in subjective effort costs cannot be fully accounted for by dopamine-related changes in the subjective weight on anticipated N-back errors.

#### Effects of Gaze and Attribute Values on Choice

As noted in the main text, gaze strongly predicted choice during the gaze-decision task (Fig. 3B): the more participants looked at either the benefits or the costs of the high-effort offer, the more likely they were to choose that offer. This result suggested that attention biases offer selection. However, we also found that it did so in an attribute-specific way. That is, gaze at hard offer benefits predicted hard offer selection more strongly than gaze at hard offer costs. This result suggested that gaze amplified the particular effect that a given attribute had on choice.

In addition, we also found that gaze predicted choice more strongly on trials in which combined offer values were higher. In a mixed effects logistic regression, for example, high-effort selection was predicted by greater proportion gaze at either feature of the high-effort offer (*β* = 0.83, P < 2.1×10^−16^), and this effect was stronger with larger summed offer values (interaction: *β* = 0.090, P = 0.0048; see Table S3 for full results). This result is consistent with the hypothesis that gaze amplified the perceived value of attended-versus-unattended offers during decision-making (*27*).

Finally, we also found evidence that gaze amplified the effect of attended versus unattended attributes in our drift diffusion modeling. Namely, our winning model contained multiplicative as well as additive terms. The simple additive effects of gaze at offer benefits (*β*_1_= 0.13, P < 2.2×10^−16^) and costs (*β*_2_ = 0.15, P < 2.2×10^−16^), indicated that gaze at either attribute predicted choice of the corresponding offer, controlling for its effect on evidence accumulation. However, we also found that the effect of benefits on choice was larger when people gazed at benefits versus costs (Eqn. 1: *β*_3_ − *β*_5_ = 0.42, P = 2.0×10^−4^). Moreover, as noted in the main text, the effect of benefits was reliably larger when gazing at benefits, while the effect of costs was larger at trend-level when gazing at costs based on pre-bifurcation gaze. These results imply that attention as indexed by gaze – and especially gaze prior to a latent choice – amplifies the effects of attended versus unattended information on the decision process.

#### Choice Reaction Times

Given the implication of striatal dopamine in vigor, we anticipated that dopamine synthesis capacity and drugs would also speed responding. Consistent with this prediction, mean response speed (inverse RT) increased on both methylphenidate and sulpiride versus placebo during both the discounting phase (Fig. S3A) and the subsequent gaze-decision task (Fig. S3B).

To ensure that this speeding was not merely a reflection of session order, or differences in characteristics of offers participants received in respective tasks, and to consider potential interactions with dopamine synthesis capacity, we regressed speed on multiple variables in fully-random hierarchical regression models. For the discounting task, we tested whether speed was predicted by session order, drug, caudate nucleus synthesis capacity, the interaction of drug and synthesis capacity, and the amount and load of the high-cost, high-benefit offer. We found that participants responded faster on methylphenidate (*β* = 0.037, P = 0.018) and sulpiride at trend-level (*β* = 0.028, P = 0.076), and moreover that the effect of sulpiride on speeding was larger for those with lower caudate dopamine synthesis capacity (drug by synthesis capacity interaction: *β* = −0.032, P = 0.0054). Additionally, participants responded faster when the base offer for the high-cost, high-benefit task was larger (€4 versus €2; *β* = 0.0088, P = 6.2×10^−4^) and in later sessions (*β*_*S*2 *vs S*1_ = 0.073, P = 1.2×10^−5^; *β*_*S*3 *vs S*1_ = 0.14, P = 2.0×10^−8^). Finally, there was a trend-level slowing when the load of the high-effort task increased (*β* = −0.0055, P = 0.092). There was neither a reliable main effect of dopamine synthesis capacity, nor an interaction between synthesis capacity and methylphenidate (both P’s ≥ 0.41).

In the subsequent gaze-decision task, we tested whether participants’ speed was influenced by the difference in offer SV (high-cost / high-benefit SV minus low-cost / low-benefit SV), absolute value differences, drug, caudate synthesis capacity, session, and trial on response speed, and found that choice difficulty and dopamine affect mean response speed across sessions. Namely, participants responded faster on easier trials (larger absolute value differences: *β* = 0.017, P = 4.5×10^−7^), and on methylphenidate versus placebo (*β* = 0.031, P = 0.038). Other significant predictors included that participants responded faster on later trials (*β* = 0.030, P = 2.9×10^−13^), in later sessions (*β*_*S*2 *vs S*1_ = 0.052, P = 0.0023; *β*_*S*3 *vs S*1_ = 0.084, P = 9.2×10^−5^), and when the subjective value of the high-cost / high-benefit option increased relative to the subjective value of the low-cost / low-benefit option (*β* = 0.0084, P = 0.0032). There was neither a reliable main effect of dopamine synthesis capacity, nor reliable interactions between drug and dopamine synthesis capacity in the caudate nucleus (all P’s ≥ 0.26).

#### Saccade Velocities

In addition to reaction times, we also measured saccade velocities to test whether striatal dopamine invigorated behavior generally. Before testing for a relationship between dopamine and saccadic velocity, however, we first regressed out the well-known relationship between saccade velocities and distances (referred to as the “main sequence”). Following (*34*), we computed the residuals of a linear slope-and-intercept model of peak velocities, in deg/s, regressed onto saccade distances in deg. Next, we averaged residual peak velocities for every trial for each participant on each drug. Variation in trial-averaged residual saccade vigor by subject and drug was consistent with the hypothesis that both methylphenidate and sulpiride induced dopamine release. In a hierarchical regression model controlling for session order, we found that sulpiride speeded saccades versus placebo (*β* = 9.87, P = 0.0021; though with no accompanying main effect of methylphenidate: *β* = 1.74, P = 0.61). Furthermore, drug by dopamine synthesis capacity interactions provided evidence that both sulpiride (*β* = - 5.72, P = 3.5×10^−4^) and methylphenidate (at trend-level: *β* = −4.48, P = 0.064) speeded saccades more for those with lower dopamine synthesis capacity. There was no main effect of dopamine synthesis capacity on residual saccade velocities (*β* = 9.17, P = 0.20).

Methylphenidate and sulpiride both increased cognitive motivation and invigorated saccades more for those with lower dopamine synthesis capacity (at trend-level for methylphenidate). If these joint effects stem from a common cause – namely, an increase in striatal dopamine release – then it is plausible that drug effects on cognitive motivation will be stronger for those who showed a greater increase in saccade velocity. To test this, we computed the mean residual saccade velocity in each drug session for each participant and then computed the difference between the mean residual velocities on methylphenidate and sulpiride versus placebo. We then fit robust linear regression models testing whether these changes also predicted drug-induced changes in the mean high-effort selection rate across participants. As shown in Fig. S4, the results of this analysis provided further evidence of increased striatal dopamine release for both drugs: the robust regression slopes were significant for sulpiride (*β* = 0.32, P = 0.013, Wald test) and trending for methylphenidate (*β* = 0.27, P = 0.074, Wald test).

While our results strongly implicate striatal dopamine, it is also possible that methylphenidate-induced changes in noradrenaline release could also explain some of our effects. In particular, noradrenaline might, to some extent, account for behavioral invigoration. However, given that sulpiride also increased vigor and that sulpiride-induced speeding predicted increased cognitive motivation, we believe that the best explanation for our collective effects involves dopamine alterations. Thus, our collective effects implicate dopamine in cognitive motivation, while not ruling out a possible complementary role for noradrenaline in behavioral vigor.

#### Drug Effect Distinctions

We anticipated that sulpiride effects might be like those of methylphenidate. This expectation was based prior work, including our own, showing that dopamine can bind pre-synaptically, in both rodents and humans, thereby increasing dopamine release (*6, 25, 35-37*). Indeed, pre-synaptic sulpiride effects been shown to amplify reward prediction error signaling and reward learning (*25*). However, sulpiride can also act post-synaptically, blocking D2 signaling (*26*). Thus, we were also open to the possibility that sulpiride effects might differ, particularly as a function of dopamine synthesis capacity.

Nevertheless, our results support the hypothesis that greater striatal dopamine release is induced by both sulpiride and methylphenidate, especially for those with lower dopamine synthesis capacity. This conclusion is buttressed by evidence that both drugs increased subjective values in the discounting task and willingness to accept the high effort offer in the gaze-decision task, in interactions with dopamine synthesis capacity. In addition, as previously noted, there was evidence that both drugs increased measures of behavioral vigor.

While methylphenidate and sulpiride had largely similar effects, they differed in that methylphenidate increased sensitivity to benefits while sulpiride decreased sensitivity to costs. This result was established by fitting logistic regressions to the gaze-decision task choice data (Eqn. S6), and was reinforced by evidence that methylphenidate further amplified the effects of benefits on evidence accumulation (Fig. 4E). The corresponding test for sulpiride revealed a non-significant suppression of the effects of costs on evidence accumulation, albeit in the expected direction. We hesitate to speculate why methylphenidate altered sensitivity to benefits while sulpiride altered sensitivity to costs. Future work should address whether this difference reflects a selective, dissociable impact of methylphenidate and sulpiride to benefit and cost dimensions, respectively, or instead reflects a shared, common mechanism (i.e. increased striatal dopamine) which could alternately impact either dimension.

#### Drug Effects on Self-Reported Affect and Medical Symptoms

Given that catecholamine drugs can impact subjective arousal, affect, and physiological symptoms, we were curious whether methylphenidate and sulpiride altered self-report measures and how these self-reported measures related to key results. We considered self-reported alertness, contentedness, calmness, PANAS positive and negative affect, and numerical ratings of various physiological symptoms including, (e.g. dizziness, headache, fatigue, etc.; collapsed to a single “medical” score). As noted, full details are provided in the on-line registration for the broader study at https://www.trialregister.nl/trial/5959.

We found no evidence that methylphenidate impacted subjective affect, arousal, or medical symptoms. None of the self-report measures were significantly different across drug sessions when measured just prior to (15 minutes before) the discounting task (all t-test P’s ≥ 0.30). Sulpiride, however, decreased negative affect (t(46) = −2.38, P = 0.021) and medical symptoms (t(46) = −2.06, P = 0.045) compared with placebo.

Next, we tested whether drug-induced changes in self-report measures predicted drug-induced changes in discounting behavior. Here we found no reliable individual difference correlations between methylphenidate-altered self-report measures and changes in AUC from the discounting task (all t-test P’s ≥ 0.17). However, sulpiride-induced increases in AUC correlated with decreases in negative affect (Spearman’s rho = −0.30, P = 0.041), increases in contentedness (rho = 0.44, P = 0.0022), and increases in alertness (rho = 0.37, P = 0.011).

We next tested whether drug-induced changes alertness, contentedness, and negative affect might explain our putative effects of dopamine on subjective values. To test this, we fit hierarchical regression models identical to Eqn. S3 to test whether subjective values were predicted by methylphenidate and sulpiride versus placebo, caudate nucleus dopamine synthesis capacity (Ki), and their interaction, but also included a single term for one of the self-report measures, and further allowed that term to vary by dopamine synthesis capacity. Thus, this test allowed us to ask whether drugs and Ki altered subjective values, controlling for self-report measures. In all cases, whether controlling for drug-induced changes in alertness, contentedness, or negative affect, sulpiride reliably interacted with Ki (all P’s ≤ 0.025). Thus, although there was shared variance, sulpiride-induced changes in affect did not explain individual differences in the effects of sulpiride on subjective value.

We also conducted parallel analyses on data from the gaze-decision task to ask whether drug-induced changes in alertness, contentedness, and negative affect could account for drug effects on sensitivity to benefit or cost information. Indeed, sulpiride-induced changes in all three measures correlated with changes in sulpiride-induced high-effort selection rates. For alertness, the correlation was (Spearman’s rho = 0.29, P = 0.047), for contentedness it was (rho = 0.45, P = 0.0019), and for negative affect it was trending (rho = −0.26, P = 0.081). No other sulpiride-altered subjective measures correlated with sulpiride-altered selection rates (all P’s ≥ 0.25). Likewise, there were no correlations between methylphenidate-altered subjective measures and methylphenidate-altered selection rates (all P’s ≥ 0.20).

To test whether self-reported alertness, contentedness, and negative affect explained the putative effects of dopamine on cost and benefit sensitivity, we fit hierarchical regression models identical to Eqn. S6, with the addition of a single main effect term for one self-report measure, and interactions between that self-report measure and benefits, costs, and Ki. These models thus allow us to test whether dopamine explains changes in benefit and cost sensitivity, controlling for changes in self-report measures. Across all our models, we found that sulpiride remained a significant (or trending in the case of alertness: P = 0.094, all other P’s ≤ 0.028) predictor of the effect of costs on choice. Thus, there was little evidence that drug-induced changes affect or mood explain drug effects on sensitivity to costs and benefits.

#### Summary Conclusions on the Effects of Early Attention

We conclude our results in the main text with the statement that: “*Collectively, our results support that striatal dopamine enhances motivation for cognitive effort by amplifying the effects of benefits versus costs attended early in a decision.*” The first part of this statement on dopamine enhancing motivation for cognitive effort is motivated by the largely consistent drug effects and their interactions with dopamine synthesis capacity (see *Drug Effect Distinctions*, above). Regarding the effects of early attention to benefits versus costs, there are multiple lines of relevant evidence. First, gaze dynamics imply that early but not late attention is most relevant for shaping choice (later gaze appears to reflect post-choice commitment). This conclusion is buttressed by evidence that pre- but not post-bifurcation gaze is multiplicative (i.e. early but not late attention amplifies the value of attended-versus-unattended offers). Second, gaze dynamics also suggest a link between choice, and attention to benefits-versus-costs, and this link appears to be strengthened by methylphenidate and increasing dopamine synthesis capacity (Fig. 3C). Third, a formal test of drift diffusion parameters revealed that higher dopamine (via dopamine synthesis capacity or methylphenidate) modulated the effects of benefits during evidence accumulation.

---

**Fig. S1.**
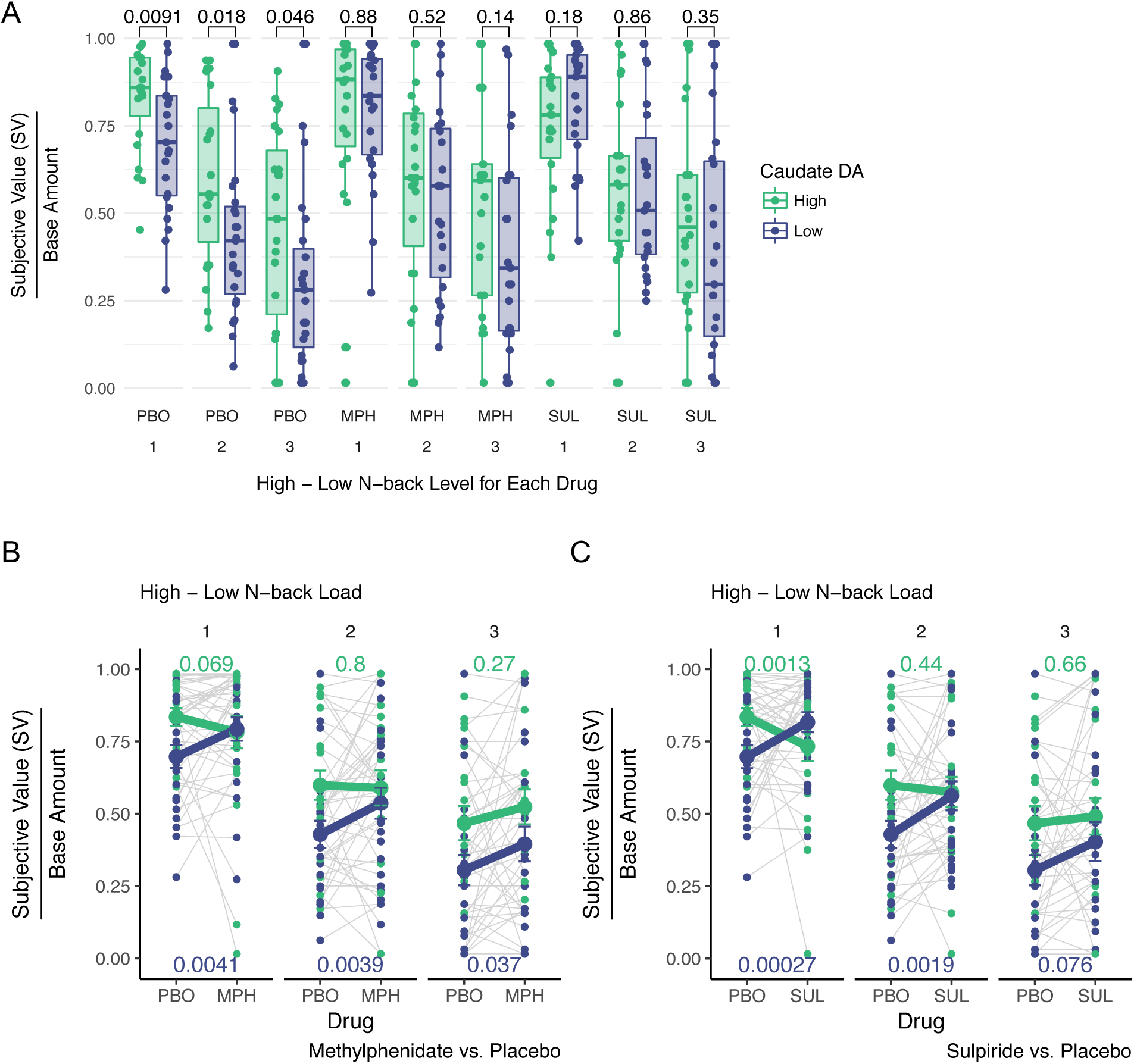
Subjective values for all participants as a function of drug, dopamine synthesis capacity in the caudate nucleus, and load differences between the high-effort and low-effort offers. **A.** Dopamine synthesis capacity separates participants’ discounting at all N-back load difference levels. P-values provided for the group comparison at every load level. In particular, for placebo, the **B.** Methylphenidate reliably increases subjective values at all load levels for participants with low dopamine synthesis capacity, but has no reliable effects for high dopamine synthesis capacity participants. P-values report results of paired, within-subjects t-tests at every load level. **C.** Sulpiride also increases subjective values at all load levels (trending for the highest level) for low synthesis capacity participants. However, while sulpiride has no effect on high synthesis capacity participants at high load level differences, it also reliably decreases subjective values for the smallest load difference among participants with high dopamine synthesis capacity.

**Fig. S2.**
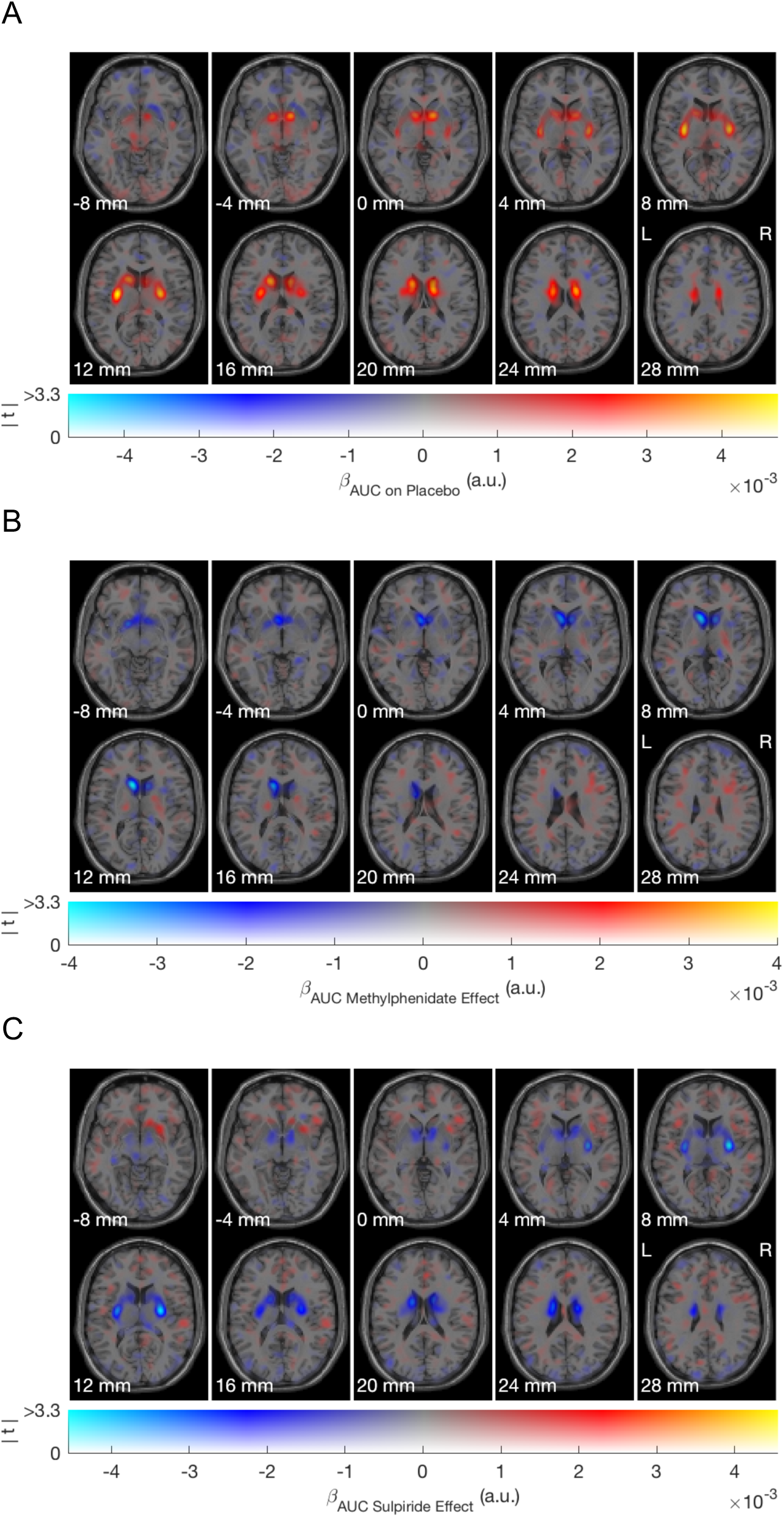
Voxel-wise dual-display of dopamine synthesis capacity (Patlak Ki) values and their interactions with drugs predicting area under the discounting curve (AUC) across individuals. Color hue represents effect size and color opacity represents t-value. Warmer and more opaque colors indicate that higher dopamine synthesis capacity predicts shallower discounting under placebo (i.e., more willingness to expend effort for reward). Dual-display figure produced using the Slice Display code from: Zandbelt, Bram (2017): Slice Display. figshare. 10.6084/m9.figshare.4742866 **A.** On placebo, shading pattern indicates a concentration in the caudate nucleus and the posterior putamen predicting AUC – though only the caudate nucleus predicted AUC reliably in our core ROI analyses. **B.** Effect of methylphenidate on AUC varies by dopamine synthesis capacity, primarily in the caudate nucleus. Here, the negative sign reflects that methylphenidate mostly increases AUC for participants with low-versus high-dopamine synthesis capacity. **C.** Effect of sulpiride on AUC varies by dopamine synthesis capacity in the caudate nucleus and in the posterior putamen. However, as noted in the Supplemental Results, the results are reliable only in the caudate nucleus: there are no reliable interactions between sulpiride versus placebo and Ki in the putamen.

**Fig. S3.**
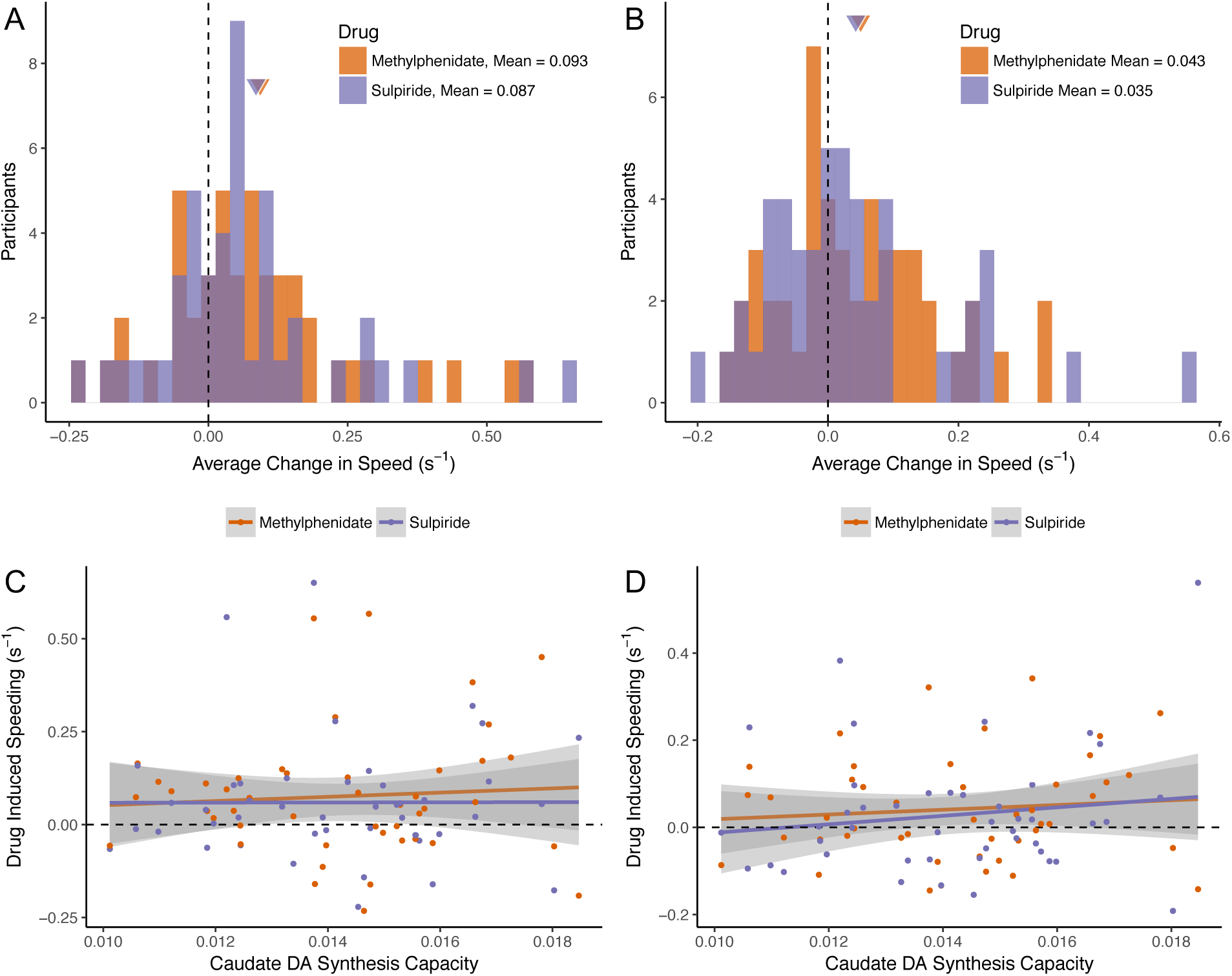
**A—B.** Drug-induced speeding as measured by inverse reaction time during the discounting task and the subsequent gaze-decision task. **C—D.** Drug induced speeding and its relationship to dopamine synthesis capacity in the caudate nucleus. **A**) During the discounting task, participants were faster on methylphenidate (P = 0.0022) and sulpiride (P = 0.0089) versus placebo, as revealed by paired t-tests of mean, inverse reaction time. **B**) During the gaze-decision task, participants were faster on methylphenidate (P = 0.014) and sulpiride at trend-level (P = 0.095). **C**) In the discounting task, there was no reliable relationship between dopamine synthesis capacity and drug-induced speeding for either drug in according to linear regression models (both P’s ≥ 0.64). **D)** In the gaze-decision task, there was no reliable relationship between dopamine synthesis capacity and drug-induced speeding for either drug in according to linear regression models (both P’s ≥ 0.64).

**Fig. S4.**
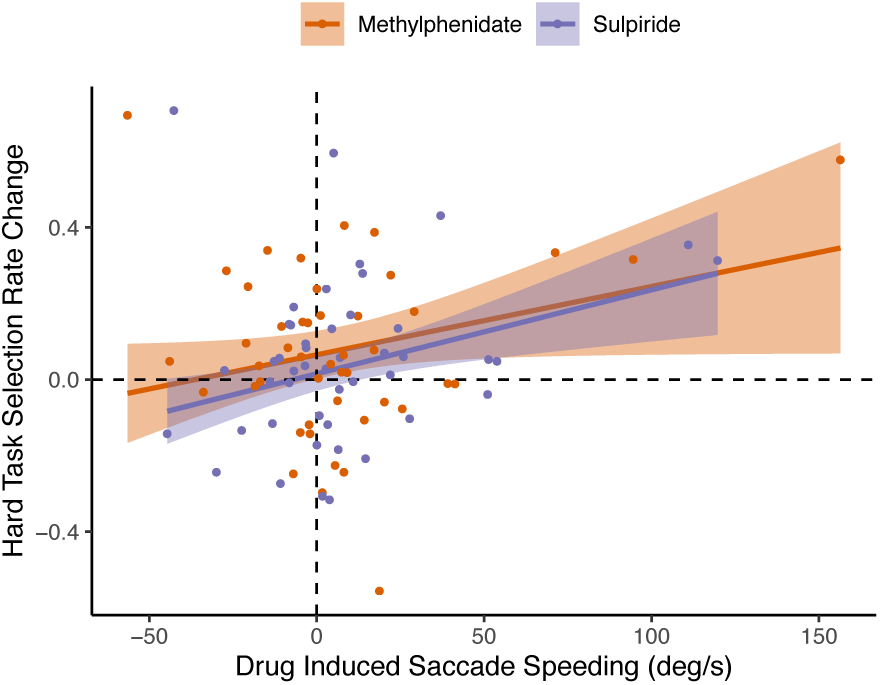
Drug-induced changes in mean residual saccade velocity predict drug-induced changes in mean hard task selection rates across participants for both sulpiride (P = 0.013, Wald test) and methylphenidate (P = 0.074, Wald test) versus placebo. Change scores computed by regressing out saccade distance, and then averaging over mean residuals by trial, then across trials by drug session. Regression lines and shading give, respectively, robust regression fits and 95% CI.

**Fig. S5.**
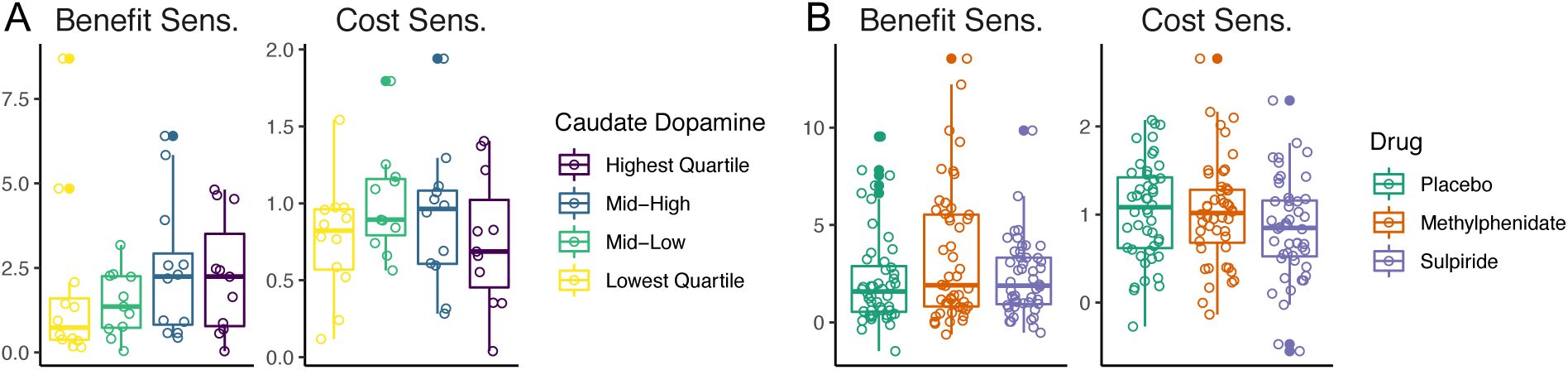
Participant-specific effect estimates (fixed plus random effects) for sensitivity to benefits and costs, as a function of dopamine synthesis capacity quartile (**A.**) or drug (**B.**). Effect estimates (change in the log-odds of choosing the high effort offer as a function of either an increase in benefits or decrease in costs) were calculated in separate models Eqns. S8—9.

**Table S1.** Table of fitted parameters of model testing effects of offer amount, drug (MPH: methylphenidate, PBO: placebo, SUL: sulpiride), dopamine synthesis capacity and relative load difference on the subjective value of high-effort offers in the discounting phase (from Eqn. S3).

| Predictor | Estimate | Standard Error | P-value |
| --- | --- | --- | --- |
| Intercept | 0.57 | 0.031 | $2.2 \times 10^{-16}$ |
| Offer Amount | 0.018 | 0.0056 | 0.0020 |
| MPH vs. PBO | 0.024 | 0.023 | 0.30 |
| SUL vs. PBO | 0.013 | 0.035 | 0.70 |
| Caudate DA | 0.068 | 0.024 | 0.0072 |
| Load Difference | -0.14 | 0.012 | $2.9 \times 10^{-15}$ |
| MPH * Caudate DA | -0.067 | 0.022 | 0.0042 |
| SUL * Caudate DA | -0.10 | 0.028 | $8.3 \times 10^{-4}$ |
| Session 2 vs. 1 | 0.0097 | 0.025 | 0.70 |
| Session 3 vs. 1 | 0.076 | 0.043 | 0.083 |

**Table S2.**
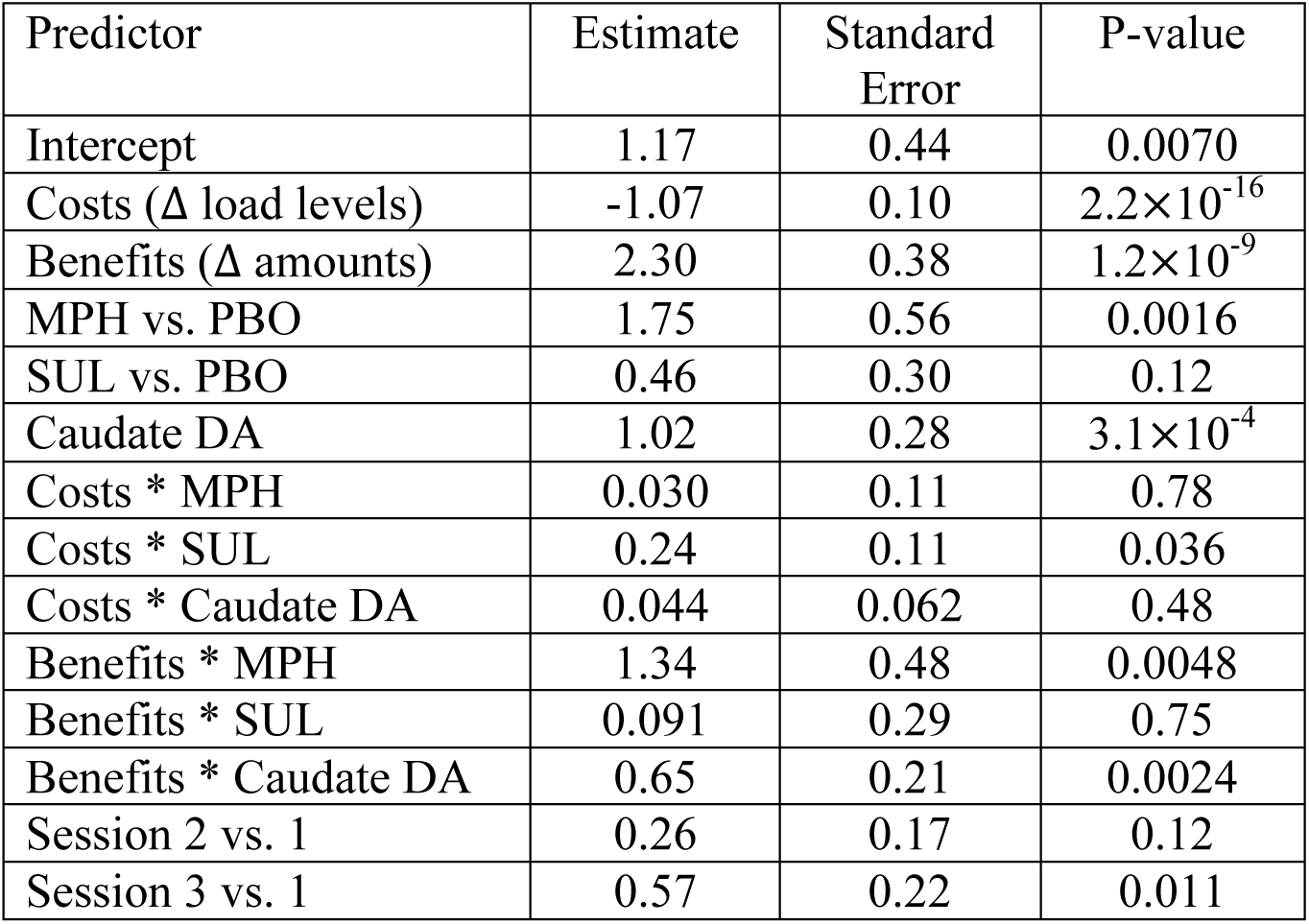
Table of fitted parameters of model testing effects of relative costs, benefits, drugs, dopamine synthesis capacity, and relevant interactions on (logistic) selection of the high-cost, high-benefit option in the gaze-decision task (from Eqn. S6).

**Table S3.** Table of fitted parameters of model testing effects of offer subjective value (SV) differences, summed SV, proportion gaze at the high-effort offer (Hi), and their interaction, as well as session number on (logistic) selection of the high-cost, high-benefit option in the gaze-decision task (from Eqn. S9).

| Predictor | Estimate | Standard Error | P-value |
| --- | --- | --- | --- |
| Intercept | 0.52 | 0.14 | $2.8 \times 10^{-4}$ |
| Hi – Lo Offer SV | 1.44 | 0.11 | $2.2 \times 10^{-16}$ |
| Summed SV | -0.0041 | 0.12 | 0.98 |
| Proportion Hi Gaze | 0.83 | 0.057 | $2.2 \times 10^{-16}$ |
| Summed SV * Prop. Hi Gaze | 0.090 | 0.032 | 0.0048 |
| Session 2 vs. 1 | 0.64 | 0.22 | 0.0036 |
| Session 3 vs. 1 | 0.32 | 0.20 | 0.12 |

**Table S4.** Table of fitted parameters of model testing effects of choice type (whether the participant selected the high-versus low-effort offer), caudate dopamine (DA) synthesis capacity, drug and their interactions on average proportion fixation of benefits versus cost information, across all time points 250—450 ms following offer onset in the gaze-decision task (from Eqn. S10).

| Predictor | Estimate | Standard Error | P-value |
| --- | --- | --- | --- |
| Intercept | -0.15 | 0.11 | 0.18 |
| Choice type (high- vs. low-effort) | 0.42 | 0.13 | 0.0017 |
| Caudate DA | -0.028 | 0.11 | 0.79 |
| MPH vs. PBO | -0.20 | 0.12 | 0.10 |
| SUL vs. PBO | -0.12 | 0.13 | 0.34 |
| Choice * Caudate DA | 0.37 | 0.13 | 0.0045 |
| Choice * MPH | 0.090 | 0.14 | 0.53 |
| Choice * SUL | 0.25 | 0.14 | 0.083 |
| MPH * Caudate DA | 0.22 | 0.12 | 0.070 |
| SUL * Caudate DA | 0.10 | 0.13 | 0.44 |
| Choice * MPH * Caudate DA | -0.36 | 0.14 | 0.012 |
| Choice * SUL * Caudate DA | -0.041 | 0.15 | 0.78 |

**Table S5.** Table of DIC values (effective number of parameters *p*_*D*_ is given in parentheses) for each model tested in HDDM using either all the data (ALL), or data from each of the individual drug sessions: placebo (PBO), methylphenidate (MPH), and sulpiride (SUL). Key model features include whether gaze and value combine additively or multiplicatively, and whether alternative offer values, or attribute values drive evidence accumulation.

| Model | Equation | All | PBO | MPH | SUL |
| --- | --- | --- | --- | --- | --- |
| Additive Net Value | S12 | 81824<br>(228) | 26822<br>(212) | 23981<br>(212) | 26060<br>(207) |
| Multiplicative Net Value | S13 | 82122<br>(229) | 26925<br>(211) | 24197<br>(212) | 26105<br>(215) |
| Additive Attributes | S14 | 79465<br>(301) | 25728<br>(275) | 22471<br>(262) | 24798<br>(244) |
| Multiplicative Attributes | S15 | 80321<br>(287) | 26124<br>(250) | 22847<br>(252) | 25068<br>(244) |
| Additive &<br>Multiplicative Net Value | S16 | 81774<br>(262) | 26814<br>(234) | 23978<br>(236) | 26034<br>(235) |
| Additive &<br>Multiplicative Attributes | 1 | 78786<br>(364) | 25577<br>(317) | 22375<br>(309) | 24657<br>(283) |

**Table S6.** Table of DIC values (effective number of parameters *p*_*D*_ is given in parentheses) for each model fit using HDDM and either pre- or post-bifurcation gaze data from either all sessions (ALL), or data from each of the individual drug sessions: placebo (PBO), methylphenidate (MPH), and sulpiride (SUL).

| Pre-bifurcation Model | Equation | All | PBO | MPH | SUL |
| --- | --- | --- | --- | --- | --- |
| Additive Attributes | S14 | 73598<br>(299) | 23684<br>(271) | 20615<br>(252) | 22703<br>(266) |
| Multiplicative Attributes | S15 | 73136<br>(300) | 23568<br>(265) | 20552<br>(261) | 22710<br>(256) |
| Additive &<br>Multiplicative Attributes | 1 | 72846<br>(370) | 23437<br>(331) | 20459<br>(308) | 22575<br>(310) |
| Post-bifurcation Model | Equation | All | PBO | MPH | SUL |
| Additive Attributes | S14 | 72811<br>(292) | 23492<br>(265) | 20369<br>(249) | 22561<br>(231) |
| Multiplicative Attributes | S15 | 73892<br>(309) | 23874<br>(269) | 20768<br>(271) | 22950<br>(268) |
| Additive &<br>Multiplicative Attributes | 1 | 72804<br>(373) | 23564<br>(323) | 20383<br>(312) | 22594<br>(294) |

**Table S7.** Table of N-back performance measure d’ mean and standard deviation, in parentheses, by N-back level and drug.

| N-back Level | PBO | MPH | SUL |
| --- | --- | --- | --- |
| 1-back | 3.55<br>(0.60) | 3.65<br>(0.60) | 3.36<br>(1.20) |
| 2-back | 3.12<br>(0.75) | 3.12<br>(0.58) | 3.25<br>(0.72) |
| 3-back | 2.00<br>(0.63) | 2.24<br>(0.85) | 2.19<br>(0.91) |
| 4-back | 1.66<br>(0.66) | 1.74<br>(0.81) | 1.75<br>(0.84) |

